# Pathologies and causes of death in stranded cetaceans in the Canary Islands (2013-2018)

**DOI:** 10.64898/2026.04.01.715953

**Authors:** Pablo Diaz-Santana, Manuel Arbelo, Josué Diaz-Delgado, Kátia Groch, Cristian Suarez-Santana, Francesco Consoli, Yara Bernaldo de Quirós, Oscar Quesada-Canales, Eva Sierra, Antonio Fernández

## Abstract

Cetacean pathology is a cornerstone for population and marine ecosystem health monitoring, allowing clear differentiation among natural and anthropogenic threats. Previous studies in the Canary Islands reported natural causes of death in 59.4% (1999-2005) and 81% (2006-2012) of stranded cetaceans, versus anthropogenic causes in 33.3% and 19%, respectively. This study aimed to determine the causes of death (CD), pathologic findings, and epidemiological patterns of 316 cetaceans stranded in the Canary Islands between 2013 and 2018. The CDs were classified in pathologic entities (PEs) emphasizing natural versus anthropic origins. Of 316 animals, 224 (70.9%) from 18 species were suitable for pathological investigations. Among natural PEE, natural pathology associated with good nutritional status (NP-GNS) and natural pathology associated with significant loss of nutritional status (NP-LNS) represented 43/224 (19.2%) and 36/224 (16%) cases, respectively. Natural pathology with undetermined nutritional status (NP-UNS) occurred in 19/224 (8.5%) animals. Intra- and interspecific traumatic interactions (ITI) represented 30/224 (13.4%) cases, followed by neonatal/perinatal pathology (NPN) 19/224 (8.5%) and live-stranding stress and/or capture myopathy (LS-CM) 18/224 (8%). Infectious and parasitic diseases predominated in natural PEs. Anthropogenic PEs included interaction with fishing activities (IFA) in 17/224 (7.6%) cases, vessel collisions (VC) in 9/22 (4%) cases, and foreign body-associated pathology (FBAP) in 3/224 (1.3%) animals. Overall, anthropogenic causes accounted for 12.9% of deaths, natural causes for 73.6%, and the CD could not be established in 30/194 (13.4%) cases. This study reaffirms the trends concerning recognized PEs (NP-GNS, NP-LNS, NP-UNS, ITI, NPN, LS-CM, IFA, VC, and FBAP), expands the body of knowledge on cetacean pathology in the Canary Islands, and reports novel findings including mixed infections, clostridiosis in uncommon species, uremic syndrome secondary to urethral nematodiasis, gas embolism in unusual species, epibiont stomatitis, congenital musculo-skeletal malformations, or neoplastic processes. These findings advance understanding of cetacean mortality patterns and support conservation and management strategies.

## Introduction

Monitoring the health status of marine mammals is a key point for conservation purposes as well as mirroring the wellness of the aquatic ecosystems [1–3]. Interestingly, cetaceans exhibit characteristics for which they are cataloged as bioindicators of the marine or marine-terrestrial interface environments (*e.g*., long life span, homeotherms, cuspid of trophic change, shore habits of many species)[1].

Diverse menaces put at risk the integrity or survival of cetaceans populations all around the world. These could be classified as anthropogenic *e.g.,* fisheries activities, pollution (acoustic, chemical, plastic), maritime traffic (*i.e*., vessel collision) and sonar usage [2–15], or natural origin (non-anthropogenic), *e.g*., biotoxins, and pathogens (viruses, bacteria, fungi, parasites)[2,3,11,15–23]. Namely pathogen pathogeneses are influenced by biotic e.g., immunological competence, genetic variability of abiotic e.g., chemical disruptors (endocrines) factors, among others [24–26].

A considerable number of microorganisms affecting cetaceans have been identified as causative agents of demolishing epizootic events (e.g., morbillivirus, influenza virus), awarded with considerable attention due to their zoonotic potential (e.g*., Brucella* spp., *Toxoplasma gondii*, *Paracoccidioidomycosis* spp.), or both [15,16,21,23,26–32].

In addition, global warming is picturing unexpected habitat and environmental changes which consequences for cetaceans are unprecedent [33,34].

As human beings are tightly linked to marine resources for subsistence, a standardized evaluation of threats and diseases affecting cetacean populations could unmask present or upcoming issues for human welfare.

Post-mortem examination of stranded cetaceans represents one of the most informative approaches for assessing population health and generating objective data to support conservation policies [6,11,24,35–38]. Recently, parallel to exponential advances in applicative artificial intelligence, the value of necropsy-derived data has been substantially enhanced through the application of machine learning tools, enabling improved data structuring and the identification of novel pathological patterns [39]

Canary Islands (Northeast Atlantic Ocean) display a series of hydrological, oceanographic, and topographic characteristics propitiating it to be one of the spots with the highest cetacean biodiversity in the world [40]. A total of 31 species, encompassing 24 odontocetes and 7 mysticetes, have been reported in the Canarian waters including resident, migratory, or seasonal species [41]. Systematic pathology-based monitoring efforts of wildlife cetacean carcasses stranded along the coast of Canary Islands have been prove to be an essential source of information for population monitoring [2,3,11]. Two previous long-lasting studies e.g., 1999-2005 and 2006-2012, assessing the causes of death in cetacean stranded in the Canary Islands, revealed and incidence of deaths caused by anthropic activities of 33% and 18%, on each period respectively. Cetacean mortality attributed to natural origin pathologies amounted to 62% and 82%, respectively [2,3]. This work aims to continue the multidisciplinary-based monitoring effort of the health status of the wildlife cetacean population within the Canary Islands, contributing to the scientific literature with new pathologic descriptions, and exposing the tendencies followed by the different pathologic entities between 2013 and 2018.

## Material and Methods

The cadavers of the stranded cetaceans along the coast of the Canary Islands, between the 1^st^ January of 2013 to the 31^st^ of December 2018, represented the main source of information for this work. No studies nor experiment with alive animals were carried out. Authorization for handling and operate with the carcasses was given by the Canary Islands Government. Stranding epidemiologic and biological data of analyzed animals were compiled (Table 1). Age category was classified following international references, focusing in gonadal development and length, encompassing the following categories: fetus/neonate/calf, juvenile/subadult, and adult [15,42–44]. The body condition (BC), based on osseous prominences (e.g., vertebrae and ribs), total mass of the epaxial musculature, blubber thickness at different levels, and considering species, age, and potential biological patterns (e.g., migration), was classified in good, moderate, poor, and emaciated [2,3]. Each carcass was also classified, depending on the decomposition condition category (DCC), as extremely fresh (individuals known to live-strand and necropsied immediately), fresh, moderate decomposition, advanced decomposition, and very advanced decomposition with mummification or skeletal remains [45].

**Table 1.**
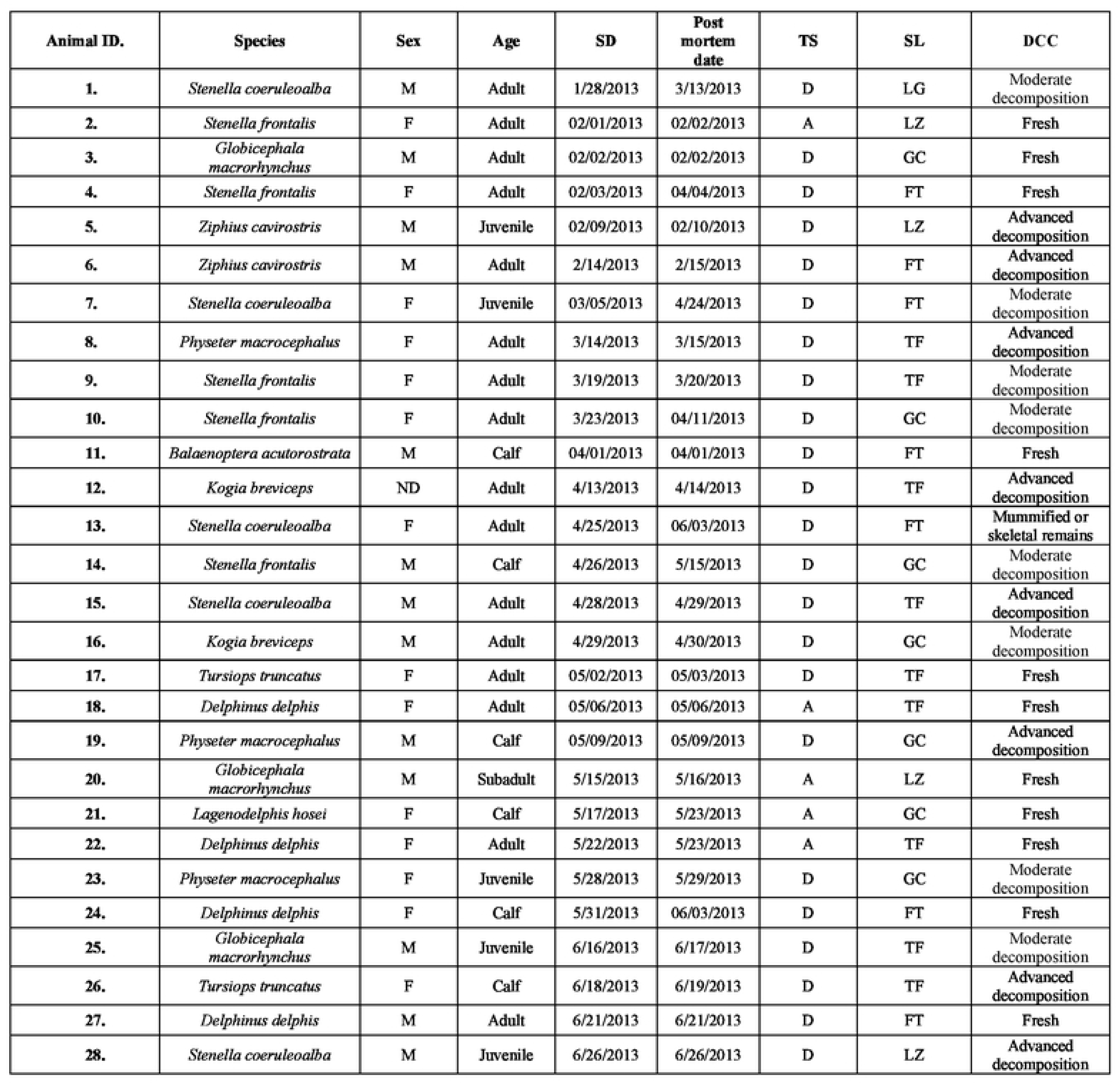
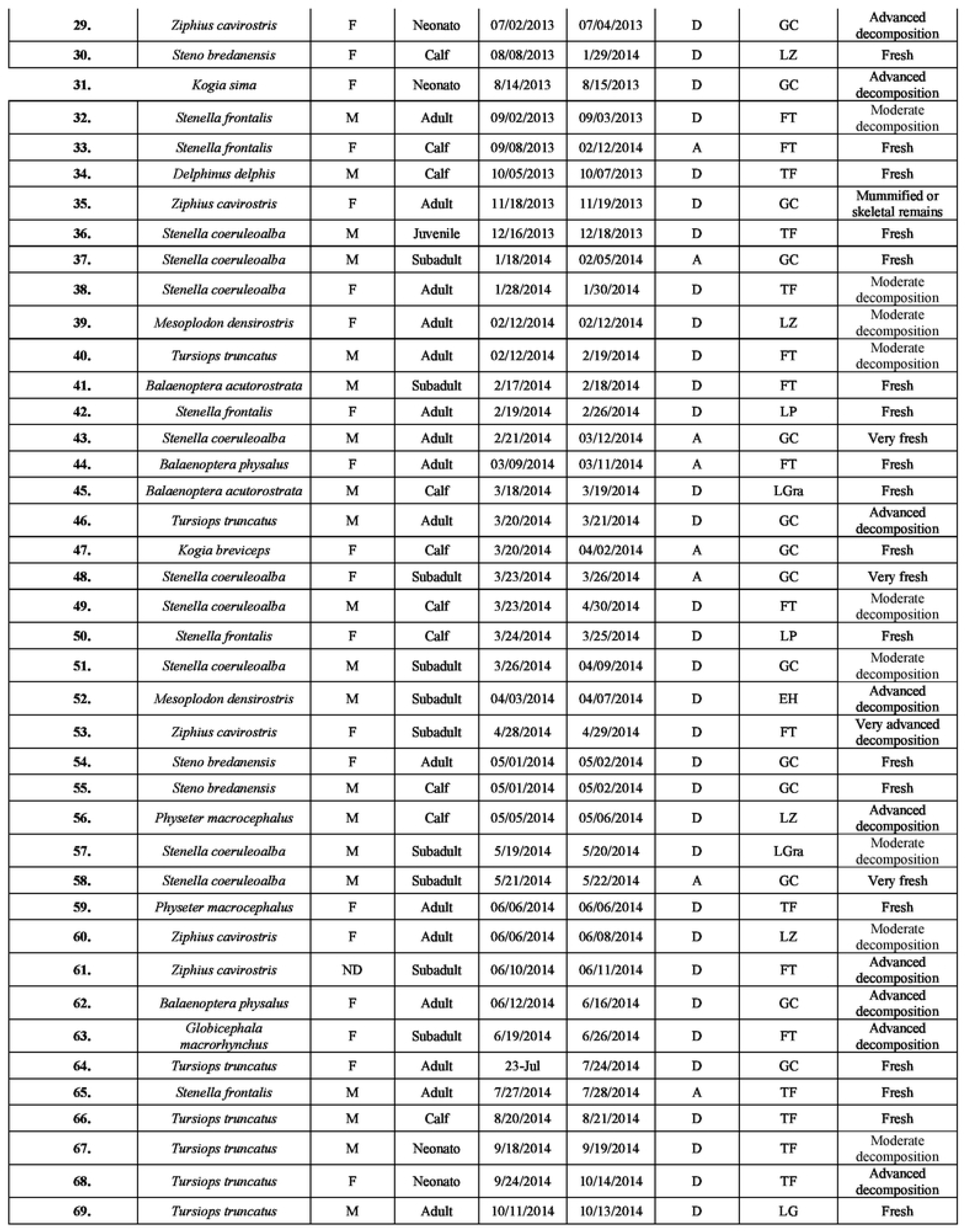
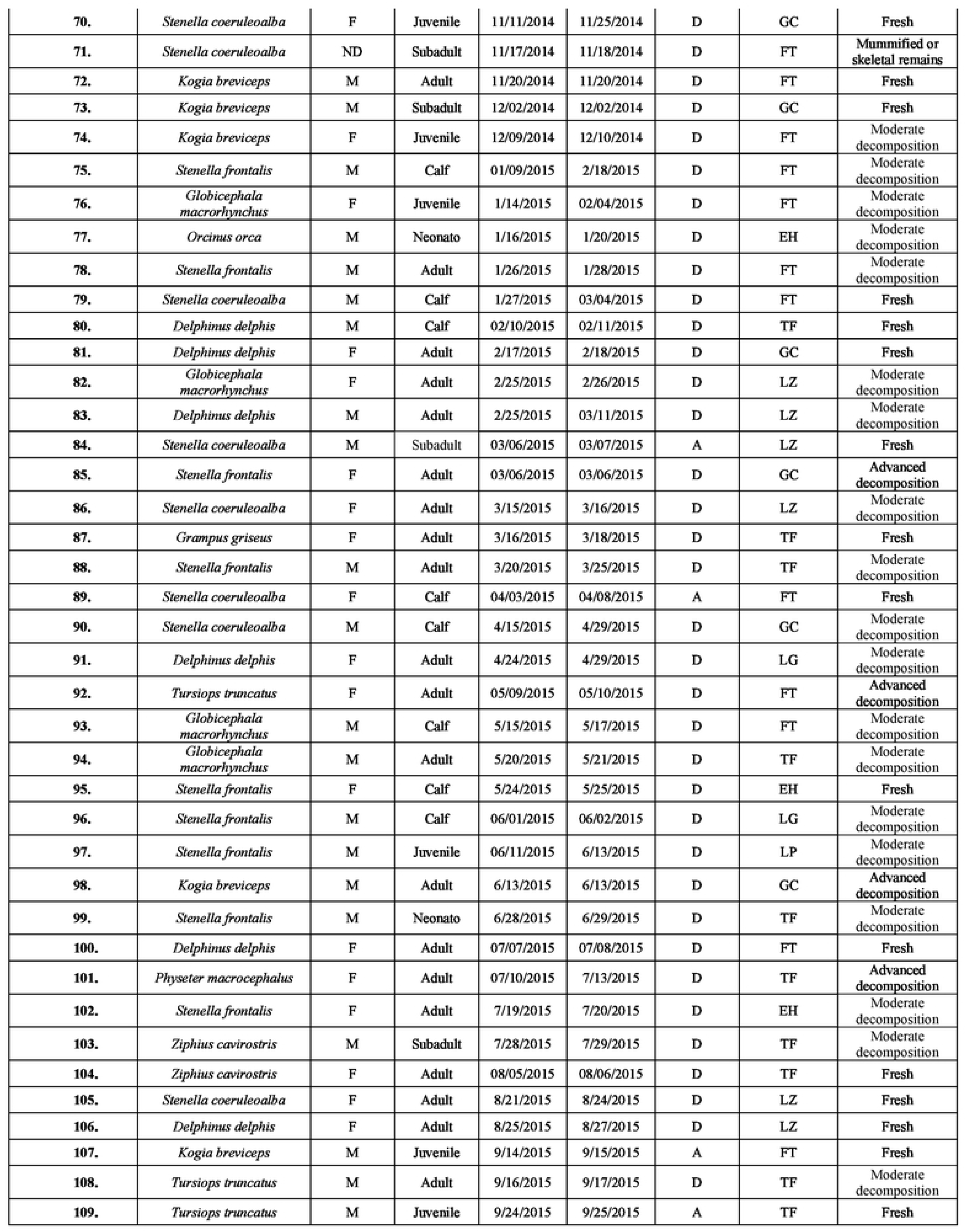
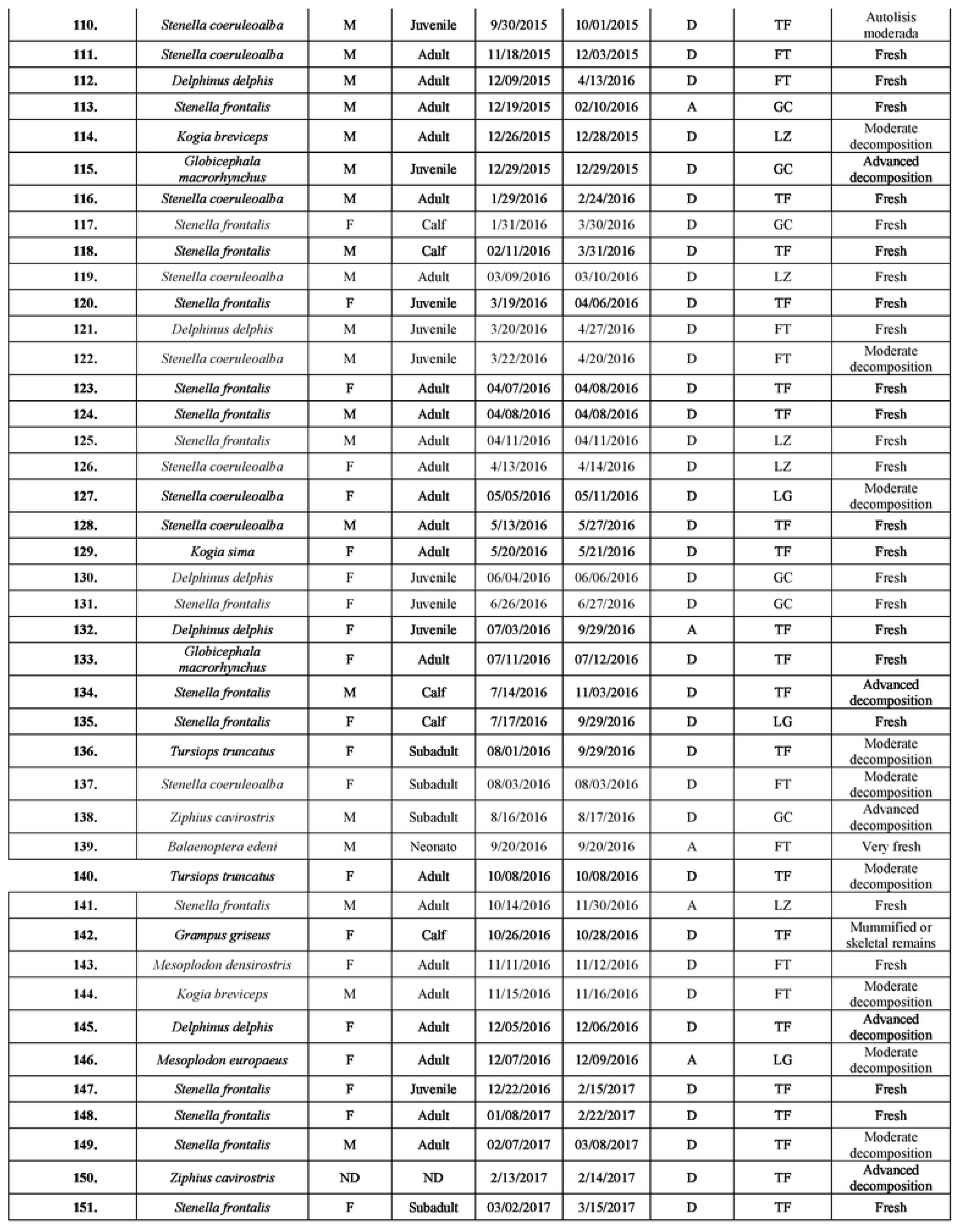
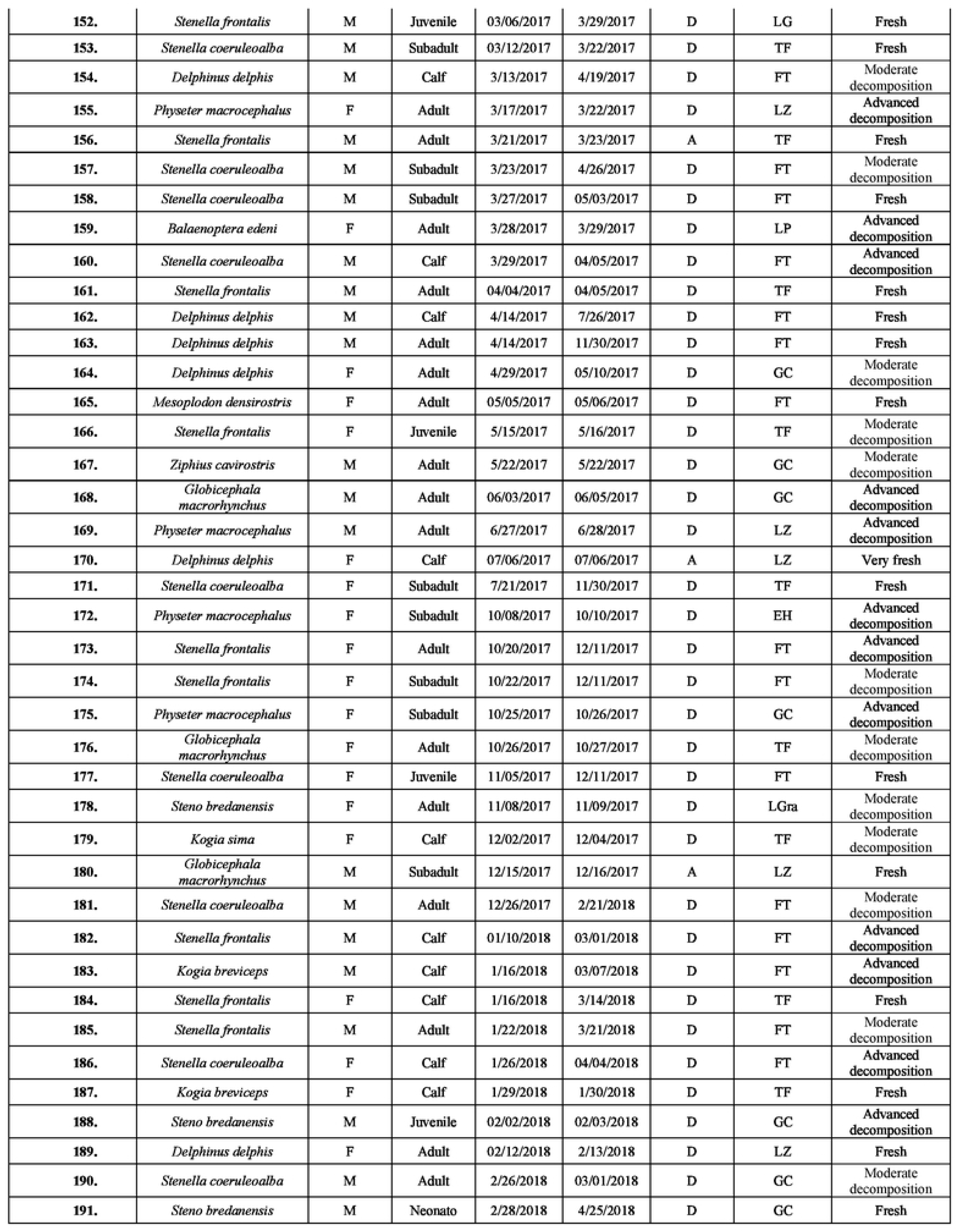
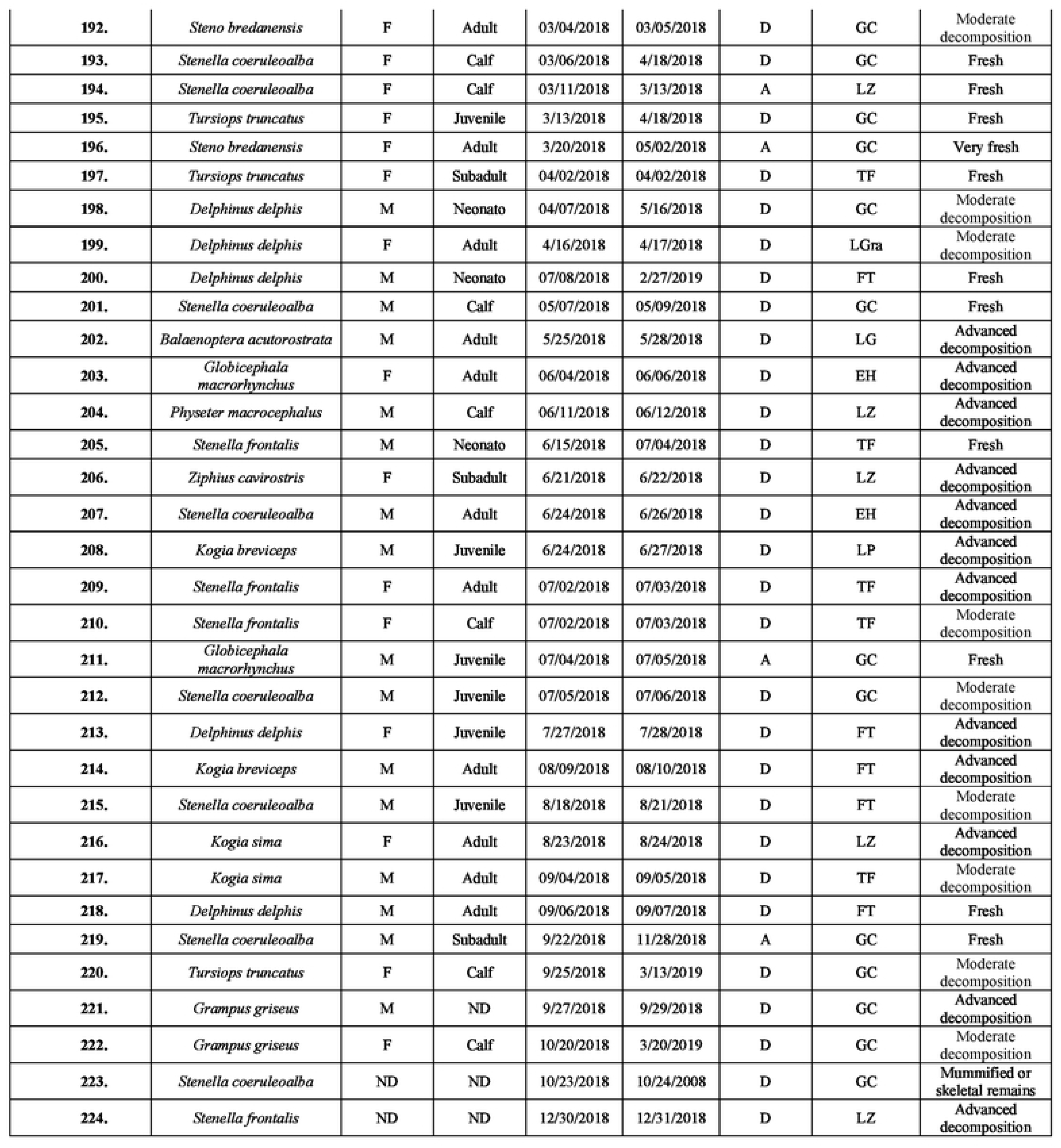
Biological and epidemiological data of 45 kogiids examined in this study. Sex: female (F), male (M), not determined (ND). Age: neonate/calf (NC),juvenile (Jv), adult (Ad). Stranding date (SD; mm/dd/yy). Type of stranding (TS; dead: D, alive: A). Stranding location (SL), island (IS): Gran Canaria (GC), Fuerteventura (FT), Lanzarote (LZ), Tenerife (TF), La Gomera (LG), El Hierro (EH), La Palma (LP), La Graciosa (LGra). Decomposition condition categories (DCC): Extremely fresh carcass (EF), fresh (F), moderate decomposition (MD), advanced decomposition (AD), mummified or skeletal remains (MSR).

| Animal ID. | Species | Sex | Age | SD | Post mortem date | TS | SL | DCC |
| --- | --- | --- | --- | --- | --- | --- | --- | --- |
| 1. | *Stenella coeruleoalba* | M | Adult | 1/28/2013 | 3/13/2013 | D | LG | Moderate decomposition |
| 2. | *Stenella frontalis* | F | Adult | 02/01/2013 | 02/02/2013 | A | LZ | Fresh |
| 3. | *Globicephala macrorhynchus* | M | Adult | 02/02/2013 | 02/02/2013 | D | GC | Fresh |
| 4. | *Stenella frontalis* | F | Adult | 02/03/2013 | 04/04/2013 | D | FT | Fresh |
| 5. | *Ziphius cavirostris* | M | Juvenile | 02/09/2013 | 02/10/2013 | D | LZ | Advanced decomposition |
| 6. | *Ziphius cavirostris* | M | Adult | 2/14/2013 | 2/15/2013 | D | FT | Advanced decomposition |
| 7. | *Stenella coeruleoalba* | F | Juvenile | 03/05/2013 | 4/24/2013 | D | FT | Moderate decomposition |
| 8. | *Physeter macrocephalus* | F | Adult | 3/14/2013 | 3/15/2013 | D | TF | Advanced decomposition |
| 9. | *Stenella frontalis* | F | Adult | 3/19/2013 | 3/20/2013 | D | TF | Moderate decomposition |
| 10. | *Stenella frontalis* | F | Adult | 3/23/2013 | 04/11/2013 | D | GC | Moderate decomposition |
| 11. | *Balaenoptera acutorostrata* | M | Calf | 04/01/2013 | 04/01/2013 | D | FT | Fresh |
| 12. | *Kogia breviceps* | ND | Adult | 4/13/2013 | 4/14/2013 | D | TF | Advanced decomposition |
| 13. | *Stenella coeruleoalba* | F | Adult | 4/25/2013 | 06/03/2013 | D | FT | Mummified or skeletal remains |
| 14. | *Stenella frontalis* | M | Calf | 4/26/2013 | 5/15/2013 | D | GC | Moderate decomposition |
| 15. | *Stenella coeruleoalba* | M | Adult | 4/28/2013 | 4/29/2013 | D | TF | Advanced decomposition |
| 16. | *Kogia breviceps* | M | Adult | 4/29/2013 | 4/30/2013 | D | GC | Moderate decomposition |
| 17. | *Tursiops truncatus* | F | Adult | 05/02/2013 | 05/03/2013 | D | TF | Fresh |
| 18. | *Delphinus delphis* | F | Adult | 05/06/2013 | 05/06/2013 | A | TF | Fresh |
| 19. | *Physeter macrocephalus* | M | Calf | 05/09/2013 | 05/09/2013 | D | GC | Advanced decomposition |
| 20. | *Globicephala macrorhynchus* | M | Subadult | 5/15/2013 | 5/16/2013 | A | LZ | Fresh |
| 21. | *Lagenodelphis hosei* | F | Calf | 5/17/2013 | 5/23/2013 | A | GC | Fresh |
| 22. | *Delphinus delphis* | F | Adult | 5/22/2013 | 5/23/2013 | A | TF | Fresh |
| 23. | *Physeter macrocephalus* | F | Juvenile | 5/28/2013 | 5/29/2013 | D | GC | Moderate decomposition |
| 24. | *Delphinus delphis* | F | Calf | 5/31/2013 | 06/03/2013 | D | FT | Fresh |
| 25. | *Globicephala macrorhynchus* | M | Juvenile | 6/16/2013 | 6/17/2013 | D | TF | Moderate decomposition |
| 26. | *Tursiops truncatus* | F | Calf | 6/18/2013 | 6/19/2013 | D | TF | Advanced decomposition |
| 27. | *Delphinus delphis* | M | Adult | 6/21/2013 | 6/21/2013 | D | FT | Fresh |
| 28. | *Stenella coeruleoalba* | M | Juvenile | 6/26/2013 | 6/26/2013 | D | LZ | Advanced decomposition |
| 29. | *Ziphius cavirostris* | F | Neonato | 07/02/2013 | 07/04/2013 | D | GC | Advanced decomposition |
| 30. | *Steno bredanensis* | F | Calf | 08/08/2013 | 1/29/2014 | D | LZ | Fresh |
| 31. | *Kogia sima* | F | Neonato | 8/14/2013 | 8/15/2013 | D | GC | Advanced decomposition |
| 32. | *Stenella frontalis* | M | Adult | 09/02/2013 | 09/03/2013 | D | FT | Moderate decomposition |
| 33. | *Stenella frontalis* | F | Calf | 09/08/2013 | 02/12/2014 | A | FT | Fresh |
| 34. | *Delphinus delphis* | M | Calf | 10/05/2013 | 10/07/2013 | D | TF | Fresh |
| 35. | *Ziphius cavirostris* | F | Adult | 11/18/2013 | 11/19/2013 | D | GC | Mummified or skeletal remains |
| 36. | *Stenella coeruleoalba* | M | Juvenile | 12/16/2013 | 12/18/2013 | D | TF | Fresh |
| 37. | *Stenella coeruleoalba* | M | Subadult | 1/18/2014 | 02/05/2014 | A | GC | Fresh |
| 38. | *Stenella coeruleoalba* | F | Adult | 1/28/2014 | 1/30/2014 | D | TF | Moderate decomposition |
| 39. | *Mesoplodon densirostris* | F | Adult | 02/12/2014 | 02/12/2014 | D | LZ | Moderate decomposition |
| 40. | *Tursiops truncatus* | M | Adult | 02/12/2014 | 2/19/2014 | D | FT | Moderate decomposition |
| 41. | *Balaenoptera acutorostrata* | M | Subadult | 2/17/2014 | 2/18/2014 | D | FT | Fresh |
| 42. | *Stenella frontalis* | F | Adult | 2/19/2014 | 2/26/2014 | D | LP | Fresh |
| 43. | *Stenella coeruleoalba* | M | Adult | 2/21/2014 | 03/12/2014 | A | GC | Very fresh |
| 44. | *Balaenoptera physalus* | F | Adult | 03/09/2014 | 03/11/2014 | A | FT | Fresh |
| 45. | *Balaenoptera acutorostrata* | M | Calf | 3/18/2014 | 3/19/2014 | D | LGra | Fresh |
| 46. | *Tursiops truncatus* | M | Adult | 3/20/2014 | 3/21/2014 | D | GC | Advanced decomposition |
| 47. | *Kogia breviceps* | F | Calf | 3/20/2014 | 04/02/2014 | A | GC | Fresh |
| 48. | *Stenella coeruleoalba* | F | Subadult | 3/23/2014 | 3/26/2014 | A | GC | Very fresh |
| 49. | *Stenella coeruleoalba* | M | Calf | 3/23/2014 | 4/30/2014 | D | FT | Moderate decomposition |
| 50. | *Stenella frontalis* | F | Calf | 3/24/2014 | 3/25/2014 | D | LP | Fresh |
| 51. | *Stenella coeruleoalba* | M | Subadult | 3/26/2014 | 04/09/2014 | D | GC | Moderate decomposition |
| 52. | *Mesoplodon densirostris* | M | Subadult | 04/03/2014 | 04/07/2014 | D | EH | Advanced decomposition |
| 53. | *Ziphius cavirostris* | F | Subadult | 4/28/2014 | 4/29/2014 | D | FT | Very advanced decomposition |
| 54. | *Steno bredanensis* | F | Adult | 05/01/2014 | 05/02/2014 | D | GC | Fresh |
| 55. | *Steno bredanensis* | M | Calf | 05/01/2014 | 05/02/2014 | D | GC | Fresh |
| 56. | *Physeter macrocephalus* | M | Calf | 05/05/2014 | 05/06/2014 | D | LZ | Advanced decomposition |
| 57. | *Stenella coeruleoalba* | M | Subadult | 5/19/2014 | 5/20/2014 | D | LGra | Moderate decomposition |
| 58. | *Stenella coeruleoalba* | M | Subadult | 5/21/2014 | 5/22/2014 | A | GC | Very fresh |
| 59. | *Physeter macrocephalus* | F | Adult | 06/06/2014 | 06/06/2014 | D | TF | Fresh |
| 60. | *Ziphius cavirostris* | F | Adult | 06/06/2014 | 06/08/2014 | D | LZ | Moderate decomposition |
| 61. | *Ziphius cavirostris* | ND | Subadult | 06/10/2014 | 06/11/2014 | D | FT | Advanced decomposition |
| 62. | *Balaenoptera physalus* | F | Adult | 06/12/2014 | 6/16/2014 | D | GC | Advanced decomposition |
| 63. | *Globicephala macrorhynchus* | F | Subadult | 6/19/2014 | 6/26/2014 | D | FT | Advanced decomposition |
| 64. | *Tursiops truncatus* | F | Adult | 23-Jul | 7/24/2014 | D | GC | Fresh |
| 65. | *Stenella frontalis* | M | Adult | 7/27/2014 | 7/28/2014 | A | TF | Fresh |
| 66. | *Tursiops truncatus* | M | Calf | 8/20/2014 | 8/21/2014 | D | TF | Fresh |
| 67. | *Tursiops truncatus* | M | Neonato | 9/18/2014 | 9/19/2014 | D | TF | Moderate decomposition |
| 68. | *Tursiops truncatus* | F | Neonato | 9/24/2014 | 10/14/2014 | D | TF | Advanced decomposition |
| 69. | *Tursiops truncatus* | M | Adult | 10/11/2014 | 10/13/2014 | D | LG | Fresh |
| 70. | *Stenella coeruleoalba* | F | Juvenile | 11/11/2014 | 11/25/2014 | D | GC | Fresh |
| 71. | *Stenella coeruleoalba* | ND | Subadult | 11/17/2014 | 11/18/2014 | D | FT | Mummified or skeletal remains |
| 72. | *Kogia breviceps* | M | Adult | 11/20/2014 | 11/20/2014 | D | FT | Fresh |
| 73. | *Kogia breviceps* | M | Subadult | 12/02/2014 | 12/02/2014 | D | GC | Fresh |
| 74. | *Kogia breviceps* | F | Juvenile | 12/09/2014 | 12/10/2014 | D | FT | Moderate decomposition |
| 75. | *Stenella frontalis* | M | Calf | 01/09/2015 | 2/18/2015 | D | FT | Moderate decomposition |
| 76. | *Globicephala macrorhynchus* | F | Juvenile | 1/14/2015 | 02/04/2015 | D | FT | Moderate decomposition |
| 77. | *Orcinus orca* | M | Neonato | 1/16/2015 | 1/20/2015 | D | EH | Moderate decomposition |
| 78. | *Stenella frontalis* | M | Adult | 1/26/2015 | 1/28/2015 | D | FT | Moderate decomposition |
| 79. | *Stenella coeruleoalba* | M | Calf | 1/27/2015 | 03/04/2015 | D | FT | Fresh |
| 80. | *Delphinus delphis* | M | Calf | 02/10/2015 | 02/11/2015 | D | TF | Fresh |
| 81. | *Delphinus delphis* | F | Adult | 2/17/2015 | 2/18/2015 | D | GC | Fresh |
| 82. | *Globicephala macrorhynchus* | F | Adult | 2/25/2015 | 2/26/2015 | D | LZ | Moderate decomposition |
| 83. | *Delphinus delphis* | M | Adult | 2/25/2015 | 03/11/2015 | D | LZ | Moderate decomposition |
| 84. | *Stenella coeruleoalba* | M | Subadult | 03/06/2015 | 03/07/2015 | A | LZ | Fresh |
| 85. | *Stenella frontalis* | F | Adult | 03/06/2015 | 03/06/2015 | D | GC | Advanced decomposition |
| 86. | *Stenella coeruleoalba* | F | Adult | 3/15/2015 | 3/16/2015 | D | LZ | Moderate decomposition |
| 87. | *Grampus griseus* | F | Adult | 3/16/2015 | 3/18/2015 | D | TF | Fresh |
| 88. | *Stenella frontalis* | M | Adult | 3/20/2015 | 3/25/2015 | D | TF | Moderate decomposition |
| 89. | *Stenella coeruleoalba* | F | Calf | 04/03/2015 | 04/08/2015 | A | FT | Fresh |
| 90. | *Stenella coeruleoalba* | M | Calf | 4/15/2015 | 4/29/2015 | D | GC | Moderate decomposition |
| 91. | *Delphinus delphis* | F | Adult | 4/24/2015 | 4/29/2015 | D | LG | Moderate decomposition |
| 92. | *Tursiops truncatus* | F | Adult | 05/09/2015 | 05/10/2015 | D | FT | Advanced decomposition |
| 93. | *Globicephala macrorhynchus* | M | Calf | 5/15/2015 | 5/17/2015 | D | FT | Moderate decomposition |
| 94. | *Globicephala macrorhynchus* | M | Adult | 5/20/2015 | 5/21/2015 | D | TF | Moderate decomposition |
| 95. | *Stenella frontalis* | F | Calf | 5/24/2015 | 5/25/2015 | D | EH | Fresh |
| 96. | *Stenella frontalis* | M | Calf | 06/01/2015 | 06/02/2015 | D | LG | Moderate decomposition |
| 97. | *Stenella frontalis* | M | Juvenile | 06/11/2015 | 6/13/2015 | D | LP | Moderate decomposition |
| 98. | *Kogia breviceps* | M | Adult | 6/13/2015 | 6/13/2015 | D | GC | Advanced decomposition |
| 99. | *Stenella frontalis* | M | Neonato | 6/28/2015 | 6/29/2015 | D | TF | Moderate decomposition |
| 100. | *Delphinus delphis* | F | Adult | 07/07/2015 | 07/08/2015 | D | FT | Fresh |
| 101. | *Physeter macrocephalus* | F | Adult | 07/10/2015 | 7/13/2015 | D | TF | Advanced decomposition |
| 102. | *Stenella frontalis* | F | Adult | 7/19/2015 | 7/20/2015 | D | EH | Moderate decomposition |
| 103. | *Ziphius cavirostris* | M | Subadult | 7/28/2015 | 7/29/2015 | D | TF | Moderate decomposition |
| 104. | *Ziphius cavirostris* | F | Adult | 08/05/2015 | 08/06/2015 | D | TF | Fresh |
| 105. | *Stenella coeruleoalba* | F | Adult | 8/21/2015 | 8/24/2015 | D | LZ | Fresh |
| 106. | *Delphinus delphis* | F | Adult | 8/25/2015 | 8/27/2015 | D | LZ | Fresh |
| 107. | *Kogia breviceps* | M | Juvenile | 9/14/2015 | 9/15/2015 | A | FT | Fresh |
| 108. | *Tursiops truncatus* | M | Adult | 9/16/2015 | 9/17/2015 | D | TF | Moderate decomposition |
| 109. | *Tursiops truncatus* | M | Juvenile | 9/24/2015 | 9/25/2015 | A | TF | Fresh |
| 110. | *Stenella coeruleoalba* | M | Juvenile | 9/30/2015 | 10/01/2015 | D | TF | Autolisis moderada |
| 111. | *Stenella coeruleoalba* | M | Adult | 11/18/2015 | 12/03/2015 | D | FT | Fresh |
| 112. | *Delphinus delphis* | M | Adult | 12/09/2015 | 4/13/2016 | D | FT | Fresh |
| 113. | *Stenella frontalis* | M | Adult | 12/19/2015 | 02/10/2016 | A | GC | Fresh |
| 114. | *Kogia breviceps* | M | Adult | 12/26/2015 | 12/28/2015 | D | LZ | Moderate decomposition |
| 115. | *Globicephala macrorhynchus* | M | Juvenile | 12/29/2015 | 12/29/2015 | D | GC | Advanced decomposition |
| 116. | *Stenella coeruleoalba* | M | Adult | 1/29/2016 | 2/24/2016 | D | TF | Fresh |
| 117. | *Stenella frontalis* | F | Calf | 1/31/2016 | 3/30/2016 | D | GC | Fresh |
| 118. | *Stenella frontalis* | M | Calf | 02/11/2016 | 3/31/2016 | D | TF | Fresh |
| 119. | *Stenella coeruleoalba* | M | Adult | 03/09/2016 | 03/10/2016 | D | LZ | Fresh |
| 120. | *Stenella frontalis* | F | Juvenile | 3/19/2016 | 04/06/2016 | D | TF | Fresh |
| 121. | *Delphinus delphis* | M | Juvenile | 3/20/2016 | 4/27/2016 | D | FT | Fresh |
| 122. | *Stenella coeruleoalba* | M | Juvenile | 3/22/2016 | 4/20/2016 | D | FT | Moderate decomposition |
| 123. | *Stenella frontalis* | F | Adult | 04/07/2016 | 04/08/2016 | D | TF | Fresh |
| 124. | *Stenella frontalis* | M | Adult | 04/08/2016 | 04/08/2016 | D | TF | Fresh |
| 125. | *Stenella frontalis* | M | Adult | 04/11/2016 | 04/11/2016 | D | LZ | Fresh |
| 126. | *Stenella coeruleoalba* | F | Adult | 4/13/2016 | 4/14/2016 | D | LZ | Fresh |
| 127. | *Stenella coeruleoalba* | F | Adult | 05/05/2016 | 05/11/2016 | D | LG | Moderate decomposition |
| 128. | *Stenella coeruleoalba* | M | Adult | 5/13/2016 | 5/27/2016 | D | TF | Fresh |
| 129. | *Kogia sima* | F | Adult | 5/20/2016 | 5/21/2016 | D | TF | Fresh |
| 130. | *Delphinus delphis* | F | Juvenile | 06/04/2016 | 06/06/2016 | D | GC | Fresh |
| 131. | *Stenella frontalis* | F | Juvenile | 6/26/2016 | 6/27/2016 | D | GC | Fresh |
| 132. | *Delphinus delphis* | F | Juvenile | 07/03/2016 | 9/29/2016 | A | TF | Fresh |
| 133. | *Globicephala macrorhynchus* | F | Adult | 07/11/2016 | 07/12/2016 | D | TF | Fresh |
| 134. | *Stenella frontalis* | M | Calf | 7/14/2016 | 11/03/2016 | D | TF | Advanced decomposition |
| 135. | *Stenella frontalis* | F | Calf | 7/17/2016 | 9/29/2016 | D | LG | Fresh |
| 136. | *Tursiops truncatus* | F | Subadult | 08/01/2016 | 9/29/2016 | D | TF | Moderate decomposition |
| 137. | *Stenella coeruleoalba* | F | Subadult | 08/03/2016 | 08/03/2016 | D | FT | Moderate decomposition |
| 138. | *Ziphius cavirostris* | M | Subadult | 8/16/2016 | 8/17/2016 | D | GC | Advanced decomposition |
| 139. | *Balaenoptera edeni* | M | Neonato | 9/20/2016 | 9/20/2016 | A | FT | Very fresh |
| 140. | *Tursiops truncatus* | F | Adult | 10/08/2016 | 10/08/2016 | D | TF | Moderate decomposition |
| 141. | *Stenella frontalis* | M | Adult | 10/14/2016 | 11/30/2016 | A | LZ | Fresh |
| 142. | *Grampus griseus* | F | Calf | 10/26/2016 | 10/28/2016 | D | TF | Mummified or skeletal remains |
| 143. | *Mesoplodon densirostris* | F | Adult | 11/11/2016 | 11/12/2016 | D | FT | Fresh |
| 144. | *Kogia breviceps* | M | Adult | 11/15/2016 | 11/16/2016 | D | FT | Moderate decomposition |
| 145. | *Delphinus delphis* | F | Adult | 12/05/2016 | 12/06/2016 | D | TF | Advanced decomposition |
| 146. | *Mesoplodon europaeus* | F | Adult | 12/07/2016 | 12/09/2016 | A | LG | Moderate decomposition |
| 147. | *Stenella frontalis* | F | Juvenile | 12/22/2016 | 2/15/2017 | D | TF | Fresh |
| 148. | *Stenella frontalis* | F | Adult | 01/08/2017 | 2/22/2017 | D | TF | Fresh |
| 149. | *Stenella frontalis* | M | Adult | 02/07/2017 | 03/08/2017 | D | TF | Moderate decomposition |
| 150. | *Ziphius cavirostris* | ND | ND | 2/13/2017 | 2/14/2017 | D | TF | Advanced decomposition |
| 151. | *Stenella frontalis* | F | Subadult | 03/02/2017 | 3/15/2017 | D | TF | Fresh |
| 152. | *Stenella frontalis* | M | Juvenile | 03/06/2017 | 3/29/2017 | D | LG | Fresh |
| 153. | *Stenella coeruleoalba* | M | Subadult | 03/12/2017 | 3/22/2017 | D | TF | Fresh |
| 154. | *Delphinus delphis* | M | Calf | 3/13/2017 | 4/19/2017 | D | FT | Moderate decomposition |
| 155. | *Physeter macrocephalus* | F | Adult | 3/17/2017 | 3/22/2017 | D | LZ | Advanced decomposition |
| 156. | *Stenella frontalis* | M | Adult | 3/21/2017 | 3/23/2017 | A | TF | Fresh |
| 157. | *Stenella coeruleoalba* | M | Subadult | 3/23/2017 | 4/26/2017 | D | FT | Moderate decomposition |
| 158. | *Stenella coeruleoalba* | M | Subadult | 3/27/2017 | 05/03/2017 | D | FT | Fresh |
| 159. | *Balaenoptera edeni* | F | Adult | 3/28/2017 | 3/29/2017 | D | LP | Advanced decomposition |
| 160. | *Stenella coeruleoalba* | M | Calf | 3/29/2017 | 04/05/2017 | D | FT | Advanced decomposition |
| 161. | *Stenella frontalis* | M | Adult | 04/04/2017 | 04/05/2017 | D | TF | Fresh |
| 162. | *Delphinus delphis* | M | Calf | 4/14/2017 | 7/26/2017 | D | FT | Fresh |
| 163. | *Delphinus delphis* | M | Adult | 4/14/2017 | 11/30/2017 | D | FT | Fresh |
| 164. | *Delphinus delphis* | F | Adult | 4/29/2017 | 05/10/2017 | D | GC | Moderate decomposition |
| 165. | *Mesoplodon densirostris* | F | Adult | 05/05/2017 | 05/06/2017 | D | FT | Fresh |
| 166. | *Stenella frontalis* | F | Juvenile | 5/15/2017 | 5/16/2017 | D | TF | Moderate decomposition |
| 167. | *Ziphius cavirostris* | M | Adult | 5/22/2017 | 5/22/2017 | D | GC | Moderate decomposition |
| 168. | *Globicephala macrorhynchus* | M | Adult | 06/03/2017 | 06/05/2017 | D | GC | Advanced decomposition |
| 169. | *Physeter macrocephalus* | M | Adult | 6/27/2017 | 6/28/2017 | D | LZ | Advanced decomposition |
| 170. | *Delphinus delphis* | F | Calf | 07/06/2017 | 07/06/2017 | A | LZ | Very fresh |
| 171. | *Stenella coeruleoalba* | F | Subadult | 7/21/2017 | 11/30/2017 | D | TF | Fresh |
| 172. | *Physeter macrocephalus* | F | Subadult | 10/08/2017 | 10/10/2017 | D | EH | Advanced decomposition |
| 173. | *Stenella frontalis* | F | Adult | 10/20/2017 | 12/11/2017 | D | FT | Advanced decomposition |
| 174. | *Stenella frontalis* | F | Subadult | 10/22/2017 | 12/11/2017 | D | FT | Moderate decomposition |
| 175. | *Physeter macrocephalus* | F | Subadult | 10/25/2017 | 10/26/2017 | D | GC | Advanced decomposition |
| 176. | *Globicephala macrorhynchus* | F | Adult | 10/26/2017 | 10/27/2017 | D | TF | Moderate decomposition |
| 177. | *Stenella coeruleoalba* | F | Juvenile | 11/05/2017 | 12/11/2017 | D | FT | Fresh |
| 178. | *Steno bredanensis* | F | Adult | 11/08/2017 | 11/09/2017 | D | LGra | Moderate decomposition |
| 179. | *Kogia sima* | F | Calf | 12/02/2017 | 12/04/2017 | D | TF | Moderate decomposition |
| 180. | *Globicephala macrorhynchus* | M | Subadult | 12/15/2017 | 12/16/2017 | A | LZ | Fresh |
| 181. | *Stenella coeruleoalba* | M | Adult | 12/26/2017 | 2/21/2018 | D | FT | Moderate decomposition |
| 182. | *Stenella frontalis* | M | Calf | 01/10/2018 | 03/01/2018 | D | FT | Advanced decomposition |
| 183. | *Kogia breviceps* | M | Calf | 1/16/2018 | 03/07/2018 | D | FT | Advanced decomposition |
| 184. | *Stenella frontalis* | F | Calf | 1/16/2018 | 3/14/2018 | D | TF | Fresh |
| 185. | *Stenella frontalis* | M | Adult | 1/22/2018 | 3/21/2018 | D | FT | Moderate decomposition |
| 186. | *Stenella coeruleoalba* | F | Calf | 1/26/2018 | 04/04/2018 | D | FT | Advanced decomposition |
| 187. | *Kogia breviceps* | F | Calf | 1/29/2018 | 1/30/2018 | D | TF | Fresh |
| 188. | *Steno bredanensis* | M | Juvenile | 02/02/2018 | 02/03/2018 | D | GC | Advanced decomposition |
| 189. | *Delphinus delphis* | F | Adult | 02/12/2018 | 2/13/2018 | D | LZ | Fresh |
| 190. | *Stenella coeruleoalba* | M | Adult | 2/26/2018 | 03/01/2018 | D | GC | Moderate decomposition |
| 191. | *Steno bredanensis* | M | Neonato | 2/28/2018 | 4/25/2018 | D | GC | Fresh |
| 192. | *Steno bredanensis* | F | Adult | 03/04/2018 | 03/05/2018 | D | GC | Moderate decomposition |
| 193. | *Stenella coeruleoalba* | F | Calf | 03/06/2018 | 4/18/2018 | D | GC | Fresh |
| 194. | *Stenella coeruleoalba* | F | Calf | 03/11/2018 | 3/13/2018 | A | LZ | Fresh |
| 195. | *Tursiops truncatus* | F | Juvenile | 3/13/2018 | 4/18/2018 | D | GC | Fresh |
| 196. | *Steno bredanensis* | F | Adult | 3/20/2018 | 05/02/2018 | A | GC | Very fresh |
| 197. | *Tursiops truncatus* | F | Subadult | 04/02/2018 | 04/02/2018 | D | TF | Fresh |
| 198. | *Delphinus delphis* | M | Neonato | 04/07/2018 | 5/16/2018 | D | GC | Moderate decomposition |
| 199. | *Delphinus delphis* | F | Adult | 4/16/2018 | 4/17/2018 | D | LGra | Moderate decomposition |
| 200. | *Delphinus delphis* | M | Neonato | 07/08/2018 | 2/27/2019 | D | FT | Fresh |
| 201. | *Stenella coeruleoalba* | M | Calf | 05/07/2018 | 05/09/2018 | D | GC | Fresh |
| 202. | *Balaenoptera acutorostrata* | M | Adult | 5/25/2018 | 5/28/2018 | D | LG | Advanced decomposition |
| 203. | *Globicephala macrorhynchus* | F | Adult | 06/04/2018 | 06/06/2018 | D | EH | Advanced decomposition |
| 204. | *Physeter macrocephalus* | M | Calf | 06/11/2018 | 06/12/2018 | D | LZ | Advanced decomposition |
| 205. | *Stenella frontalis* | M | Neonato | 6/15/2018 | 07/04/2018 | D | TF | Fresh |
| 206. | *Ziphius cavirostris* | F | Subadult | 6/21/2018 | 6/22/2018 | D | LZ | Advanced decomposition |
| 207. | *Stenella coeruleoalba* | M | Adult | 6/24/2018 | 6/26/2018 | D | EH | Advanced decomposition |
| 208. | *Kogia breviceps* | M | Juvenile | 6/24/2018 | 6/27/2018 | D | LP | Advanced decomposition |
| 209. | *Stenella frontalis* | F | Adult | 07/02/2018 | 07/03/2018 | D | TF | Advanced decomposition |
| 210. | *Stenella frontalis* | F | Calf | 07/02/2018 | 07/03/2018 | D | TF | Moderate decomposition |
| 211. | *Globicephala macrorhynchus* | M | Juvenile | 07/04/2018 | 07/05/2018 | A | GC | Fresh |
| 212. | *Stenella coeruleoalba* | M | Juvenile | 07/05/2018 | 07/06/2018 | D | GC | Moderate decomposition |
| 213. | *Delphinus delphis* | F | Juvenile | 7/27/2018 | 7/28/2018 | D | FT | Advanced decomposition |
| 214. | *Kogia breviceps* | M | Adult | 08/09/2018 | 08/10/2018 | D | FT | Advanced decomposition |
| 215. | *Stenella coeruleoalba* | M | Juvenile | 8/18/2018 | 8/21/2018 | D | FT | Moderate decomposition |
| 216. | *Kogia sima* | F | Adult | 8/23/2018 | 8/24/2018 | D | LZ | Advanced decomposition |
| 217. | *Kogia sima* | M | Adult | 09/04/2018 | 09/05/2018 | D | TF | Moderate decomposition |
| 218. | *Delphinus delphis* | M | Adult | 09/06/2018 | 09/07/2018 | D | FT | Fresh |
| 219. | *Stenella coeruleoalba* | M | Subadult | 9/22/2018 | 11/28/2018 | A | GC | Fresh |
| 220. | *Tursiops truncatus* | F | Calf | 9/25/2018 | 3/13/2019 | D | GC | Moderate decomposition |
| 221. | *Grampus griseus* | M | ND | 9/27/2018 | 9/29/2018 | D | GC | Advanced decomposition |
| 222. | *Grampus griseus* | F | Calf | 10/20/2018 | 3/20/2019 | D | GC | Moderate decomposition |
| 223. | *Stenella coeruleoalba* | ND | ND | 10/23/2018 | 10/24/2008 | D | GC | Mummified or skeletal remains |
| 224. | *Stenella frontalis* | ND | ND | 12/30/2018 | 12/31/2018 | D | LZ | Advanced decomposition |

Necropsies were performed following internationally accepted protocols [44–46]. Samples from skin, blubber, *longissimus dorsi* and *rectus abdominis*, tongue, teeth, esophagus, larynx and larynx tonsil, oro-pharynx and pharynx tonsil, thyroid, thymus, trachea, bronchi, lung, heart, pericardium, aorta (thoracic and abdominal), rete mirabile, diaphragm, stomach chambers (keratinized, glandular, pyloric), intestine, pancreas, spleen, liver, kidneys, urinary bladder, adrenal glands, lymph nodes (prescapular, tracheobronchial, pulmonary, mesenteric, retroperitoneal), testes and ovaries, uterus and vagina, penis, mammary gland, cerebrum, cerebellum, encephalic trunk, VIII cranial nerve, spinal cord, hypophysis, eyes, pterygoid and nasal sinuses, mandibular and melon acoustic fat, and tympano-periotic complexes were collected and fixed in 10% neutral buffered formalin. Tissues were processed routinely, embedded in paraffin-wax and 5 μm-thick sections were stained with hematoxylin and eosin (H&E) for histologic analysis.

Tissue sections (3-10 μm) for ancillary histochemical techniques were performed when considered, including: periodic acid-Schiff, luxol fast blue, Prussian blue, , Gram/Twort, Hall’s, Grocott-Gomori’s Methenamine Silver, Masson’s trichrome, Movat-Russel pentachromat, osmium tetroxide (post-fixation), chromic acid (frozen samples), rhodamine, Congo red, von Kossa, and Ziehl-Neelsen [47,48].

Following histologic features and aiming to identify suspicious infectious agents (i.e., cetacean morbillivirus [CeMV], Herpesvirus [Hv], *Erysipelothrix rhusiopathiae*, *Brucella ceti*) immunohistochemistry (IHC) was performed in specific cases following published and standardized protocols [49–52]. Negative controls consisted of tissue sections in which a non-immune homologous serum was used instead of the primary antibodies. Positive controls encompassed well-preserved and immunoreactive cetacean, canine and/or human tissues, respectively. Further IHC information can be consulted in Table 2.

**Table 2.** Details of the primary antibodies used in this study, including information on the provider, clonality, dilution, pretreatment, incubation, secondary antibody used for each primary antibody, its source, and the detection system employed. CDV: Canine Distemper Virus. HSV1: Human Herpes Simplex Virus type 1. HIER: Heat-Induced Epitope Retrieval. ABC: Avidin-Biotin-Peroxidase Complex. ^*1^ Antibody generated from a strain isolated from an animal infected with *E. rhusiopathiae* from a previous study period (Díaz-Delgado et al. 2015). ^*2^ Source: Portanti et al. 2006, Di Febo et al. 2012.

| Primary antibody | Source/provider | Clonality | Dilution | Pretreatment | Incubation | Secondary antibody | Source/provider | Detection |
| --- | --- | --- | --- | --- | --- | --- | --- | --- |
| Morbillivirus | VMRD | Monoclonal CDV-N | 1:100 | HIER (autoclave) | 18 h, 4°C | Rabbit anti-mouse | DAKO | ABC |
| *Erysipelothrix rhusiopathiae* | Non-commercial antibody*<sup>1</sup> | Polyclonal | 1:100 | Pronasa 10' | 18 h, 4°C | Rabbit anti-mouse | DAKO | ABC |
| Herpesvirus | Abcam | Polyclonal HSV1 | 1:200 | HIER (autoclave) | 18 h, 4°C | Pig anti-mouse | DAKO | ABC |
| *Toxoplasma gondii* | VMRD | Polyclonal 7. gondii antisera | 1:400 | HIER (citrate) | 18 h, 4°C | Rabbit anti-goat | DAKO | ABC |
| *Brucella* sp. | Institute Zooprofilattico Sperimentale dell'Abruzzo e del Molise Giuseppe Caporale | Monoclonal (4B5A) | 1:10 / 1:100 | HIER (citrate) | 8', 120°C | Mouse anti-pig | In-house *<sup>2</sup> | ABC |

Molecular diagnostic techniques, particularly PCR-based methods and when considered after histologic evaluation, were employed to confirm the presence of specific pathogens. DNA/RNA was extracted from 300 µm of tissue using pressure filtration with the QuickGene Mini 80 and DNA Tissue Kit S, with RNA carrier added during lysis [53]. CeMV was detected by one-step RT-PCR targeting a 426 bp region of the P gene, nested RT-PCR for the P gene (35), and real-time RT-PCR targeting a 192 bp region of the F gene [53–55]. Hv DNA was identified using nested PCR targeting the DNA polymerase gene [56]. *Brucella* spp. were identified using duplex qPCR targeting the IS711 gene and PCR amplifying a 223 bp fragment of the *bcsp31* gene [57,58]. *T. gondii* was detected by real-time PCR targeting a 529 bp repeat and a 163 bp 18S rRNA region (GenBank AY663792) using Primer3-designed primers. *Nasitrema* spp. was identified via primers targeting a 230 bp NADH dehydrogenase gene fragment (GenBank KT180216).

For bacteriological and toxicological analyses, tissue samples (skin, muscle, lung, liver, kidney, spleen, lymph nodes, intestine, brain) were preserved at –80 °C. Analyses were conducted at VISAVET (Madrid) and IUSA (ULPGC). Bacteriologic results are recorded in Table 3. In decomposed specimens, etiological interpretation was approached with caution. Toxicology data are part of separate, unpublished studies.

**Table 3.**
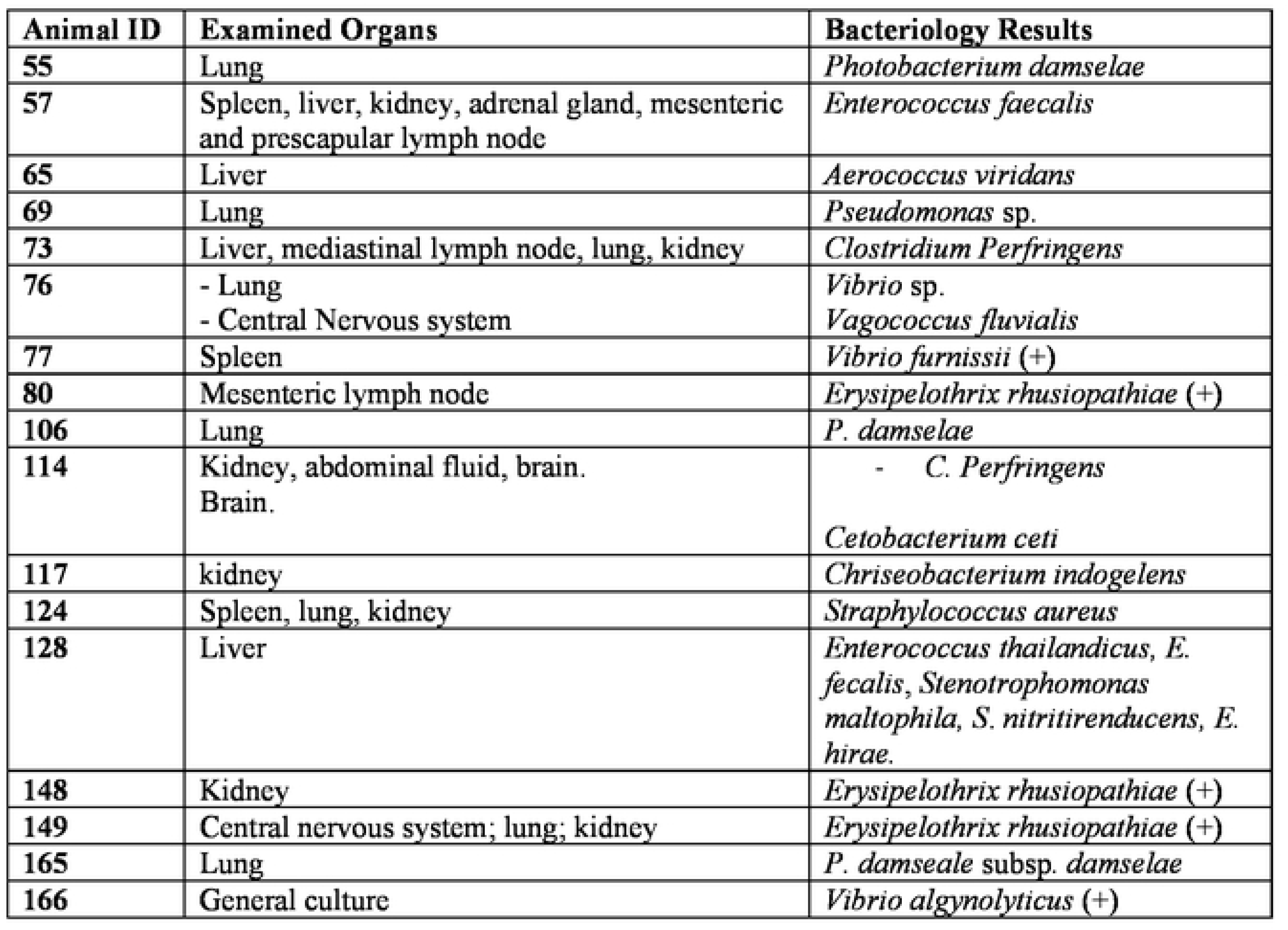
Bacterial isolations in 224 stranded cetaceans in the Canary Islands (2013-2018).

Ecto- and endoparasites, including epibionts, were collected during necropsy and preserved in 70% ethanol. Identification was based on morphological features, with PCR used for select groups (e.g., trematodes, nematodes) with related data published elsewhere [59,60].

When possible, imaging analysis (*i.e*., CT scans) of whole or specific anatomical parts was performed using a Helical CT (Toshina Astellion 16) at 120 kV, 300 mA, with 1 mm slices and 0.8 mm intervals. Reconstructions used standard and high-resolution bone algorithms.

Following predecessor long-term studies and aiming homogenization of data, the main classification method focused on differentiation of natural and anthropogenic CCD [2,3]. Several subclassifications (*i.e*., pathologic entities) were implemented in an effort to refine the most plausible cause of death within this groups, including: natural pathology associated with good nutritional status (NPGNS; including good and moderate animals), natural pathology associated with significant loss of nutritional status (NPSLNS; including poor and emaciated animals), natural pathology without stablished nutritional status (NPWNS; including animals without established body condition), intra- and interspecific traumatic interactions (ITI), live-stranding stress and /or capture myopathy-related pathology (LSSP), and neonatal/perinatal pathology (NPP) within natural CCD. Comprising anthropogenic CD, we defined the following: interaction with fishing activities (IFA), vessel collision (VC), and foreign body-associated pathology (FBAP) [2,3].

## Results

### 1.1. Stranding epidemiology

A total of 316 cetacean strandings were recorded along the coasts of the Canary Islands between January 1^st^, 2013, and December 31^st^, 2018. Of these, postmortem examinations were conducted in 224 individuals (71%), representing 18 species. A total of 215 odontocete (15 species) and 9 mysticeti (3 species) were analyzed. During this period, the average number of strandings and necropsies per year was 53 and 37, respectively, with stranding peaks occurring between The majority of strandings were registered in the central and eastern islands (*i.e*., Tenerife, Gran Canaria, Fuerteventura, and Lanzarote). Analyzed animals were 107 (47.76%) females and 111 (49.55%) males; the sex could not be determined in 6 cases (2.67%). The age categories were: 103 (45.98%) adults, 61 (27.23%) subadults/juveniles, and 57 (25.44%) neonates/calves; the age could not be determined in 3 animals. Cadaver decomposition status was very fresh in 6 (2.67%), fresh in 85 (37.94%), moderate decomposition in 56 (25%), advanced decomposition in 70 (31.25%), and very advanced decomposition in 7 (3.12%) animals. Nutritional status distribution encompassed: 11 (4.91%) good, 70 (31.25%) moderate, 59 (26.33%) poor, and 6 (2.67%) very poor/cachectic. In 78 (34.82%) animals, the nutritional status could not be assessed reliably. From 224 animals, 29 (12.9%) stranded alive and 195 (87.1%) stranded dead or were first sought floating offshore.

The most probable CD, classified in PE, was diagnosed in 194/224 (86.6%) animals (Table 4).

**Table 4.** Pathologic categories and etiologic entities in 224 stranded cetaceans the Canary Islands (2013-2018). Af: number of affected animals, Ev: number of evaluated

| Cause of death | Pathologic entities | Cases | Af/Ev (%) |
| --- | --- | --- | --- |
| *Natural or Non-anthropogenic* | Pathology not associated with significant loss of nutritional status | 1, 2, 4, 14, 20, 27, 37, 39, 40, 44, 48, 50, 54, 70, 72, 80, 83, 87, 93, 95, 97, 105, 109, 111, 116, 120, 121, 123, 124, 125, 136, 141, 147, 148, 149, 158, 163, 171, 177, 181, 184, 215, 219 | 43/224<br>(19.2%) |
|  | Pathology not associated with significant loss of nutritional status | 3, 7, 10, 17, 23, 49, 55, 58, 64, 65, 66, 69, 73, 76, 86, 100, 106, 112, 114, 117, 118, 119, 128, 131, 132, 143, 162, 164, 165, 166, 174, 176, 192, 197, 202, 206, 212 | 36/224<br>(16%) |
|  | Natural origin pathology without assessable nutritional status | 6, 15, 36, 51, 57, 61, 68, 78, 90, 92, 96, 102, 103, 150, 190, 204, 209, 210 | 19/224<br>(8.5%) |
|  | Neonatal/perinatal pathology | 24, 26, 29, 31, 45, 67, 75, 77, 99, 134, 135, 139, 170, 191, 193, 198, 200, 205, 222 | 19/224<br>(8.5%) |
|  | Traumatic intra-interspecific interaction | 8, 9, 13, 16, 38, 25, 74, 82, 91, 94, 98, 127, 129, 133, 144, 146, 154, 157, 160, 167, 175, 179, 185, 186, 188, 195, 203, 208, 216, 217 | 30/224<br>(13.4%) |
|  | Live-stranding stress and /or capture myopathy-related pathology | 18, 21, 22, 33, 41, 43, 84, 89, 107, 178, 180, 187, 189, 194, 196, 199, 201, 211 | 18/224<br>(8%) |
| *Anthropogenic* | Interaction with fishing activities | 11, 32, 34, 42, 81, 88, 113, 115, 122, 130, 145, 151, 152, 153, 156, 161, 218 | 17/224<br>(7.6%) |
|  | Foreign-body associated pathology | 5, 47, 79 | 3/224<br>(1.3%) |
|  | Vessel collision | 13, 20, 54, 60, 102, 105, 139, 160, 215 | 9/22<br>(4%) |
| *Undetermined* |  | 28, 30, 35, 46, 52, 56, 60, 62, 63, 71, 85, 108, 110, 126, 137, 140, 142, 155, 168, 169, 172, 173, 182, 183, 207, 213, 220, 221, 223, 224 | 30/194<br>(13.4%) |

### 1.2. Pathology not associated with significant loss of nutritional status (NP-GNS)

GNS was diagnosed in 43/224 (22.16%) animals. Etiologic diagnoses included: infectious (58.13%), parasitic (27.9%), neoplastic (2.3 %), gas embolism (2,32%), and others (9.3%; intestinal perforations, non-infectious cerebral necrosis, dystocia with pyometra).

Among viral pathogens, CeMV was identified by IHC and/or PCR in two short-finned pilot whales (animals no. 20 and 93) and one striped dolphin (*S. coeruleoalba*, animal no. 219). These animals had lymphoplasmacytic encephalitis with perivascular cuffing, gliosis, neuronal necrosis, lymphoplasmacytic interstitial pneumonia with type II pneumocyte hyperplasia, lymphoid depletion, necrotizing pharyngeal and laryngeal tonsilitis, and occasional viral inclusion bodies. One short-finned pilot whale had concomitant cerebral mucormycosis. Herpesvirus (HV) infection was confirmed by IHC and/or PCR in a striped dolphin (animal no. 105), a pygmy sperm whale (*Kogia breviceps*, animal no. 734), and two Atlantic spotted dolphins (*S. frontalis*, animals no. 124 and 184). These animals had lymphoplasmacytic meningoencephalitis with perivascular cuffing, gliosis, neuronal necrosis, and hypophysitis (animals no. 72, 105, and 124), as well as dermatitis (animal no. 72), all with frequent intranuclear viral inclusion bodies. Confirmed (IHC and/or PCR) concomitant infectious diseases were septicemia by *Staphylococcus aureus* (no. 124) (Fig. 1A) and *Brucella* spp. (no. 184). Cutaneous poxvirosis was diagnosed in a Risso’s dolphin (*Grampus griseus*; animal no. 87). Inflammatory central nervous system (CNS) infection of undetermined etiology was noted in 12/43 (27.9%) animals.

**Fig. 1.**
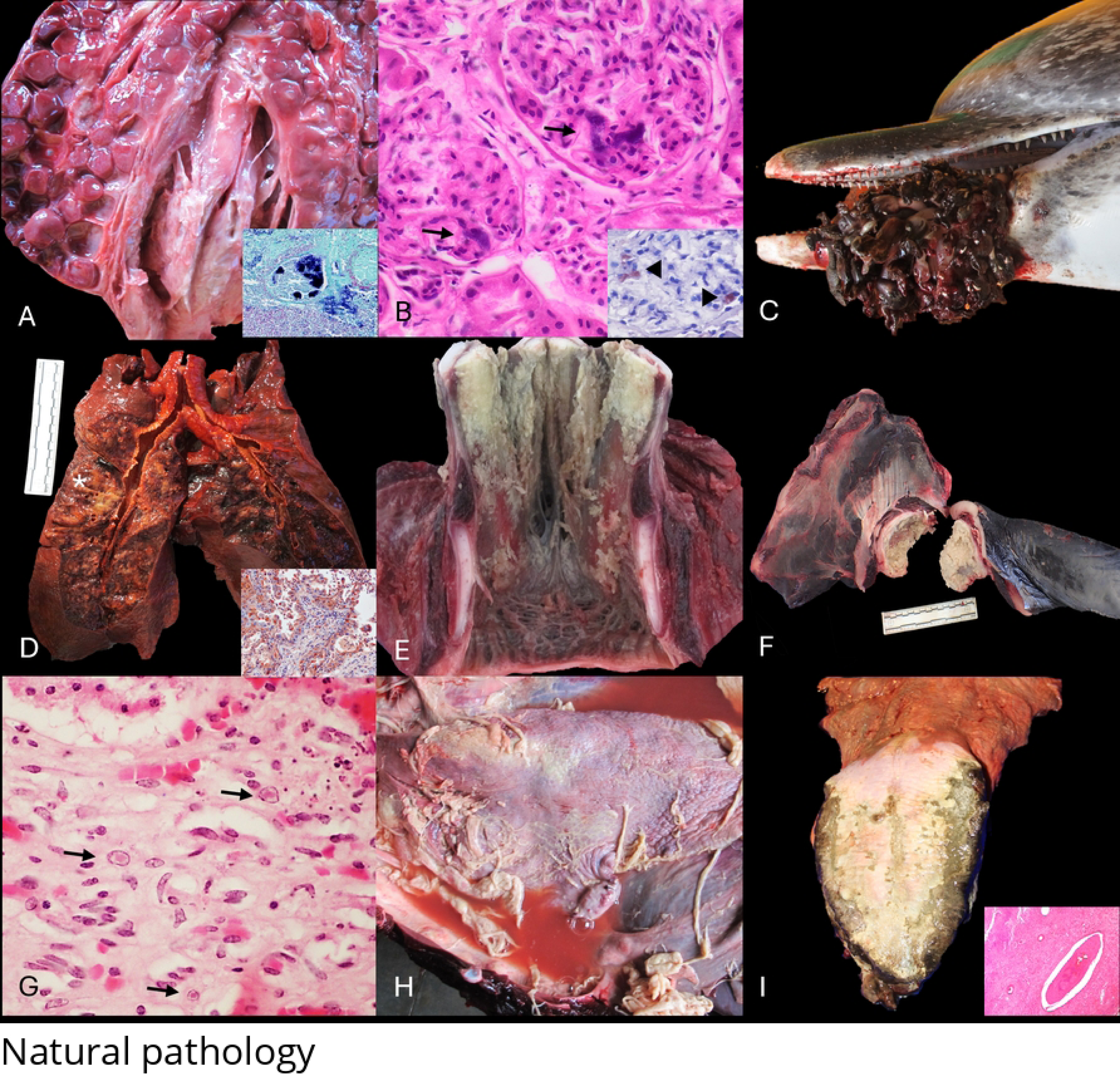
Panel of natural origin pathologies in cetaceans stranded in the Canary Islands (2013-2018). **A.** *Staphylococcus aureus* infection (animal no. 124; *S. frontalis*). Marked, diffuse necrosuppurative pyelonephritis and ureteritis. Inset: Detail of an intravascular coronary vessel thrombi with abundant embedded gram-positive bacteria. Gram stain. **B.** *Erysipelothrix rhusiopathiae* infection (animal no. 148; *S. frontalis)*. Multifocal bacterial emboli are lodged within glomerular tuft capillaries (black arrows). H&E. Inset: multifocal positive immunolabeling for *E. rhusiopathiae* within the glomerular tuft (black arrow heads). IHC for *E. rhusiopathiae*. **C.** Epibiontic stomatitis (barnacle-associated) (animal no. 123; *S. frontalis*). Focally extensive and unilateral necroulcerative and proliferative stomatitis with exposition of the left mandibular bone (not visible) and abundant intralesional cirripeds (*Conchoderma auritum*) causing severe malocclusion. **D.** Morbillivirus infection (animal no. 132; *Delphinus delphis*). Marked, multifocal to coalescing interstitial pneumonia with multifocal areas of parenchymal consolidation (asterisk). Morbillivirus infection (animal no. 132; *D. delphis*). Bronchial epithelium and multinucleated giant cells exhibiting strong cytoplasmic positive immunolabelling. IHC for morbillivirus. **E.** Morbillivirus infection (animal no. 76; *Globicephala macrorynchus*). Longitudinal view of the larynx (goose beak). Marked, multifocal to coalescence fibrinonecrotizing suppurative laryngitis. **F.** Brucella sp. infection (animal no. 165; *Mesoplodon densirostris*). Right scapulo-humeral joint (glenoid fossa and humeral head). Marked, diffuse fibrinonecrotizing arthritis. **G.** Herpesviral infection (animal no. 165; *M. densirostris)*. Marked, multifocal interstitial lymphoplasmacytic nephritis with necrosis of tubuloepithelial cells and numerous, intranuclear, 4-6 µ, eosinophilic inclusion bodies within renal tubular epithelium (black arrows). H&E **H.** *Pseudomonas* sp. infection (animal no. 69; *Tursiops truncatus*). Marked, diffuse fibrinous pleurobronchopneumonia. **I.** Uremic syndrome due to urethral obstruction by *Crassicauda* sp. (Animal no. 3; *G. macrorhyncus)*. Marked, bilateral necroulcerative glossitis. Inset: Detail of intravascular thrombi in a medium caliber artery whitin the tongue. H&E.

Fatal bacterial infections included septicemia by *Brucella* spp. in a striped dolphin (animal no. 70) and an Atlantic spotted dolphin (animal no. 184) [61], and septicemia by *E. rhusiopathiae* in two Atlantic spotted dolphins (animals nos. 148 and 149) (Fig. 1B). and one short-beaked common dolphin (*Delphinus delphis*, animal no. 80).

Severe parasitic infections involved pterygoid sinusitis by *Nasitrema* spp., *Crassicauda* spp. and *Stenurus* spp., often concomitantly, in 6 animals from 5 species (animals nos. 1, 4, 27, 87, 148). Severe sarcocystosis was diagnosed in one short-beaked common dolphin (animal no. 83), one striped dolphin (animal no. 111), one Atlantic spotted dolphin (animal no. 797), and one Risso’s dolphin (animal no. 87). Multisystemic crassicaudiasis often involving epaxial and hypaxial myositis and fasciitis was seen in 12/43 (27.9%). Prostatic crassicaudiasis was seen in one short-beaked common dolphin (animal no. 793) and one Atlantic spotted dolphin (animal no. 822).An Atlantic spotted dolphin had necroulcerative stomatitis with numerous *Conchoderma auritum* maxillopods (animal no. 123) (Fig. 1C).

Neoplastic disease was noted in an Atlantic spotted dolphin (S. frontalis, animal no. 2) with a uterine leiomyoma [62], and concurrent fibrinonecrotizing endometritis and postpartum uterine prolapse Fatal systemic gas embolism was diagnosed in a Blainville beaked whale (*Mesoplodon densirostris*; animal no 39). The animal presented multisystemic intravascular gas bubbles and a remarkably dilated right auricle and aortic bulb. Multisystemic intravascular gas bubbles associated with acute perivascular hemorrhage were confirmed; notably, the CNS was severely affected.

### 1.3. Pathology not associated with significant loss of nutritional status (NP-LNS)

NP-LNS included 36/ 224 (16%) animals. Etiologic diagnoses included: infectious (61.1%), parasitic (30.5%), and others (8.3%; kypho-lordosis , pneumothorax, polycystic gas hepatopathy).

Among viral causes, CeMV was identified in one short-finned pilot whale (*Globicephala macrorhynchus*, no. 76), one short-beaked common dolphin (no. 132), and one striped dolphin (no. 212). These animals exhibited typical inflammatory lesions in the CNS, respiratory system, and multisystemic lymphoid depletion (Fig. 1D). An uncommon lesion was fibrinonecrotizing laryngitis with squamous metaplasia in the pilot whale included, which also had suppurative bronchopneumonia by *Vibrio* spp (Fig. 1E).

HV infection was confirmed in seven odontocetes representing five species; three of them (animals no. 76, 132, 212) had concurrent morbillivirosis. CNS and pulmonary inflammatory changes were common in these animals.

A Blainville’s beaked whale (animal no. 165) had arthritis and pneumonia associated with *Brucella* sp. and *Photobacterium damselae* subsp. *damselae*, and HV-nephritis (Fig. 1F and 1G). HV-nephritis was also detected in another Blainville’s beaked whale (animal no. 143) that had concurrent pyogranulomatous encephalitis by *Nasitrema delphini* [63]. *Pseudomonas* sp. was associated to fibrinosuppurative pleurobronchopneumonia and pyothorax in an Atlantic bottle-nose dolphin (animal no. 69; Fig. 1H). *Photobacterium damselae* was ascribed to severe suppurative bronchopneumonia and septicemia in two rough-toothed dolphins (*Steno bredanensis*, mother and calf, animals nos. 55, 106). Septicemia by *Cl. Perfringens* was identified in two pygmy sperm whales (animals nos. 73 and 114). Five other animals (four odontocetes and one mysticete), had histologic evidence of bacterial infection including orchitis (animal no. 65) and endocarditis (animal no. 17); however, their etiologies remain unknown.

An Atlantic spotted dolphin (animal no. 131) had pyogranulomatous tracheitis, thyroiditis, and laryngitis by *Rhizopus arrhizus* as the etiologic agent [64].

A short-finned pilot whale (animal no. 3) had severe urethral obstruction due to *Crassicauda* and concurrent pyelonephritis along with necroulcerative glossitis, tracheitis, and laryngitis, and multiorgan thrombosis (Fig. 1I).

A short-beaked common dolphin (animal no. 112) had a right-sided pneumo(hemi)thorax. Severe intraluminal nematodal burden (genus/species) resulted in local rupture of the right tracheal bronchus . Severe congenital lumbar kypho-lordosis was identified in an Atlantic spotted dolphin calf (animal no 118). Radiographic image analysis revealed the presence of hemivertebrae in the affected segment.

”Polycystic gas hepatopathy” recapitulating features of Budd-Chiari-like syndrome was diagnosed in a striped dolphin (animal no. 128)[65].

A cavernous hemangioma was identified in a mesenteric lymph node of a short-beaked common dolphin (animal no. 132).

### 1.4. Natural origin pathology without assessable nutritional status (NPWNS)

NPWNS accounted for 19/224 (8.9%) animals from 5 species of odontocetes. Etiologic diagnoses included: parasitic (61.1%), infectious (27.7%), and others (dystocia, metabolic steatohepatitis; 11.11%).

Five Cuvier’s beaked whales (animals no. 6, 61, 103, 150, 206) had moderate to severe endarteritis arteritis by *Crassicauda* spp. larvae and renal parasitism by adult stages. The severity of vascular and renal lesions correlated with the age of the animal.

Among viral diseases, CeMV was confirmed in a striped dolphin (animal no. 51) with lymphoplasmacytic meningoencephalitis, polyradiculoneuriitis, interstitial pneumonia, and lymphoid depletion.

Severe atlanto-occipital arthritis and marked fibrinosuppurative balanoposthitis were seen in a striped dolphin (animal no. 15) and a sperm whale (animal no. 204), respectively. The suspected underlying infectious etiologies remained undetermined.

An Atlantic spotted dolphin (animal no. 102) had laceration of the left uterine horn and suppurative peritonitis secondary to dystocia. The fetus exhibited marked macrosomia with mandibular brachygnathia, subcutaneous edema, erupted teeth, and multiple vertebral fractures.

Metabolic steatohepatitis was determined in a striped dolphin (animal no. 36) on the basis of, gross hepatomegaly and icterus, and histologic features of hepatocellular vacuolar degeneration, hepatocellular regenerative nodules, cholestasis and periportal hepatitis.

### 1.5. Neonatal/perinatal pathology (NPN)

NPN was identified in 19/224 animals (8.5%), comprising 11 species (9 odontocetes and 2 mysticetes). Etiologic diagnoses included fetal distress (63.1%), dystocia (26.3%), premature maternal-social separation and/or neonatal weakness (15.7%), intra- or interspecific trauma (15.7%), and infectious causes (15.7%).

Animals with evidence of fetal distress had pulmonary edema with foamy macrophages and abundant intra-alveolar keratin squames, as well as atelectasis and emphysema. Tracheal and pulmonary (aspirated) meconium was noted in a short-beaked common dolphin neonate (animal no. 198; Fig. 2A). Alveolar hyaline membranes and severe oedema were observed in animals nos. 26 and 67.

**Fig. 2.**
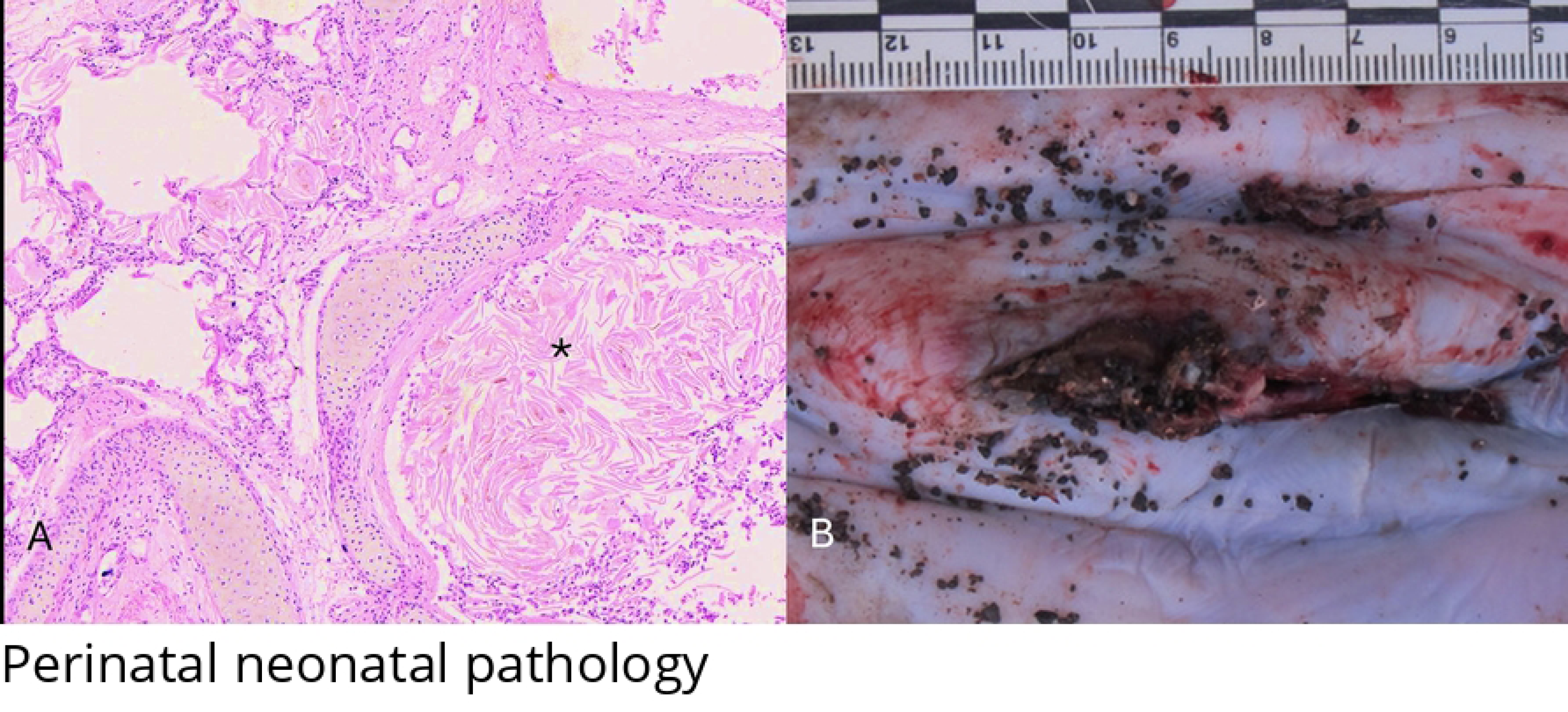
Panel of neonatal/ perinatal pathologies in cetaceans stranded in the Canary Islands (2013-2018). **A.** Pulmonary aspiration (animal no. 198; *D. delphis*). Multiple bronchi and alveoli are filled with abundant keratin squames (asterisk) and sloughed epithelial cells. **B.** Umbilical patency (animal no. 45; *Balaenoptera edeni*). Swollen and erythematous umbilicus.

A killer whale fetus (*Orcinus orca*; animal no. 77) had evidence of fetal distress, suppurative hepatitis and adrenalitis with necrosis and intralesional bacteria. Bacteriologic analysis identified *Vibrio furnisii. A* Bryde’s whale (*Balaenoptera edeni*; animal no 819) had suppurative omphalitis and lymphadenitis, and cardiac hemorrhage (Fig. 2B). The suspected infectious cause could not be determined.

### 1.6. Traumatic intra-interspecific interaction (IITI)

IITI was determined in 30/224 (13.4%) including 10 species of odontocetes. The striped dolphin and the short-finned pilot whale were overrepresented.

The main gross findings in these cases were cavitary and multiorgan hemorrhage, most prominently affecting the lungs, bone fractures (e.g., ribs, vertebrae, mandible, and scapula), pulmonary perforations, skull fractures, and cutaneous “tooth-rake” lesions were common (Fig. 3A and 3B). Histologically, myonecrosis with discoid degeneration was prevalent. Pigmentary tubulonephrosis was identified in five animals (nos. 16, 82, 94, 175, 203), representing four deep-diving species. Additional findings were tracheal and pulmonary edema, pulmonary emphysema, and partially digested gastric content.

**Fig. 3.**
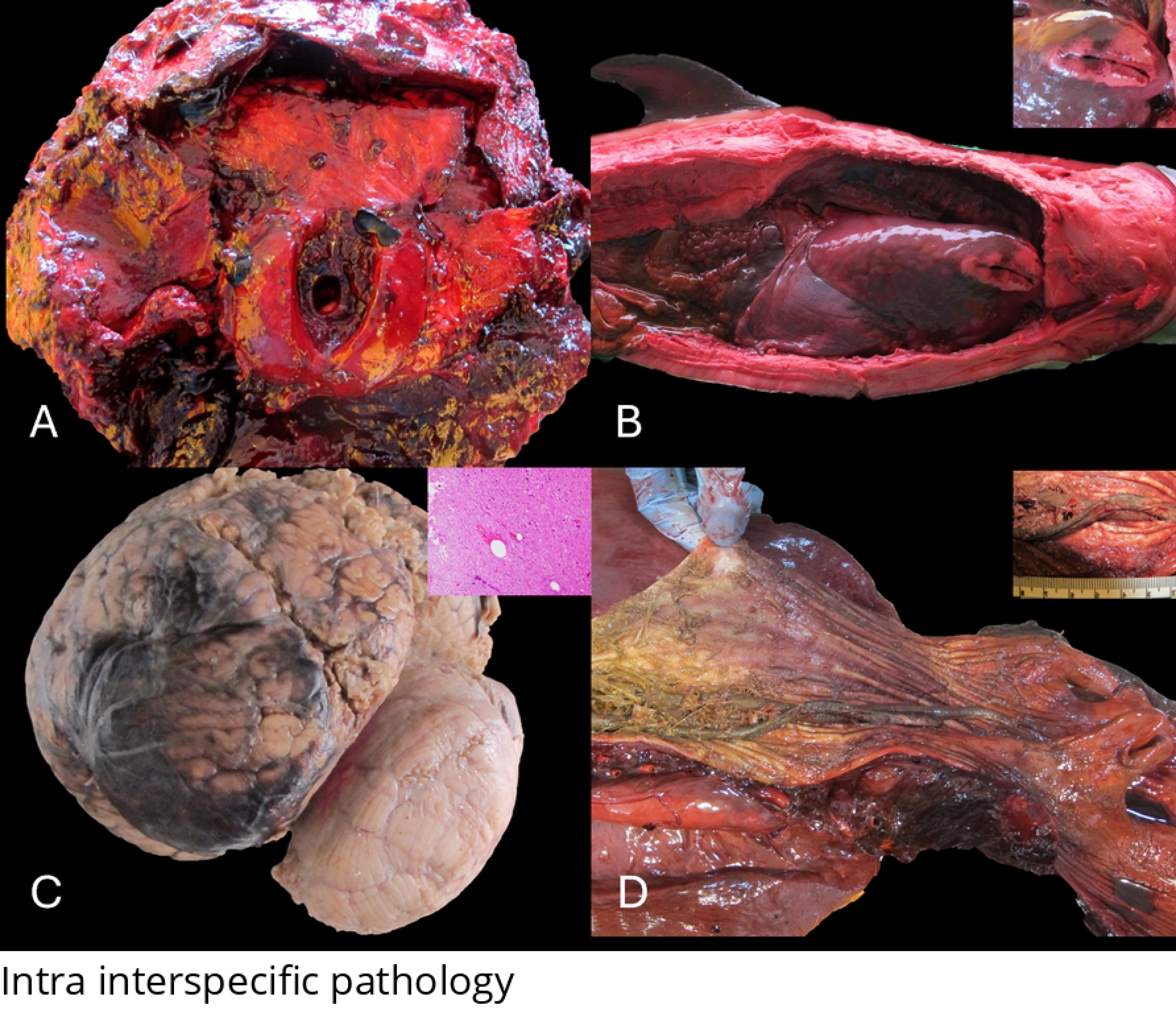
Panel of pathologies associated to intra and interspecific interactions in cetaceans stranded in the Canary Islands (2013-2018). **A.** Neurocraneum fractures and hemorrhages (animal no. 129; *Kogia sima*). Occipital bone fracture with cervical and subdural hemorrhages (not visible). **B.** Cerebral hemorrhage (animal no. 133; *G. macrorhyncus*). Focally extensive hemorrhage within the left frontal lobe following trauma. Bottom left inset: Detail of gas bubbles expanding the neuropile with peripheral radiated hemorrhages. **C.** Lung perforation (animal no. 9; *S. frontalis*). Right lung with focal cranio-dorsal perforation. Inset: Detail of perforation with multifocal hemorrhages, marginal alveoli hyperinflation and elevated retracted borders. **D.** Fatal predator prey interaction (animal no. 195; *T. truncatus*). Longitudinal section of the esophagus with one intraluminal, twisted , 30-40 cm, poorly digested eel (morphologically compatible with *Conger conger*) causing deviation of the larynx towards the nasopharyngeal funnel (not visible). Inset: Detail of the *C. conger* with diffuse grey coloration and multifocal yellow spots.

A short-finned pilot whale (animal no. 133) had cranioencephalic trauma and multisystemic gas embolism with severe involvement of the CNS (Fig. 3C), and a Sowerby’s beaked whale (*Mesoplodon bidens*, animal no. 146) with multiorgan intravascular myofiber embolism following an IITI [66].

A total of five individuals (animal nos. 74, 91, 185, 216, 217) presented cutaneous rake marks compatible with killer whale interactions, occasionally accompanied by massive soft and bone tissue loss.

Relevant co-morbidities were identified in several IITI animals, including herpesviral necro-hemorrhagic and ulcerative dermatitis in a pygmy sperm whale (animal no. 74), morbillivirosis and partial penile urethral obstruction and pyogranulomatous nephritis in a short finned pilot whale (animal no. 94), lymphoplasmacytic myelitis and myocarditis in an Atlantic spotted dolphin (animal no. 9), lymphoplasmacytic encephalitis with perivascular cuffing in a striped dolphin (animal no. 157), and a Cuvier’s beaked whale (animal no. 167) with *Crassicauda* spp. arteritis and pulmonary herpesviral infection. A bottlenosed dolphin (animal no. 195) had laryngeal luxation and aspiration pneumonia associated with two elongated fish, morphologically compatible with congers (*Conger conger*) (Fig. 3D).

### 1.7. Live-stranding stress and /or capture myopathy-related pathology (LSS/CMP)

LSS/CMP involved 18/224 (8%) individuals belonging to 8 species (7 odontocetes; 1 mysticete). Typical gross lesions were subcutaneous and muscular hemorrhage and necrosis, concomitant with cutaneous lacerations primarily in the rostral, ventral, and cranial regions of the pectoral and caudal fins (Fig. 4A and 4B). Histologically, acute rhabdomyolysis with contraction band necrosis were common in hypoaxial, epaxial, and cardiac muscle, as well as multiorgan hemorrhage, pigmentary tubulonephrosis (Fig. 4C), and cytoplasmic hepatocellular hyaline globules, and alveolar, tracheal, and laryngeal edema.

**Fig. 4.**
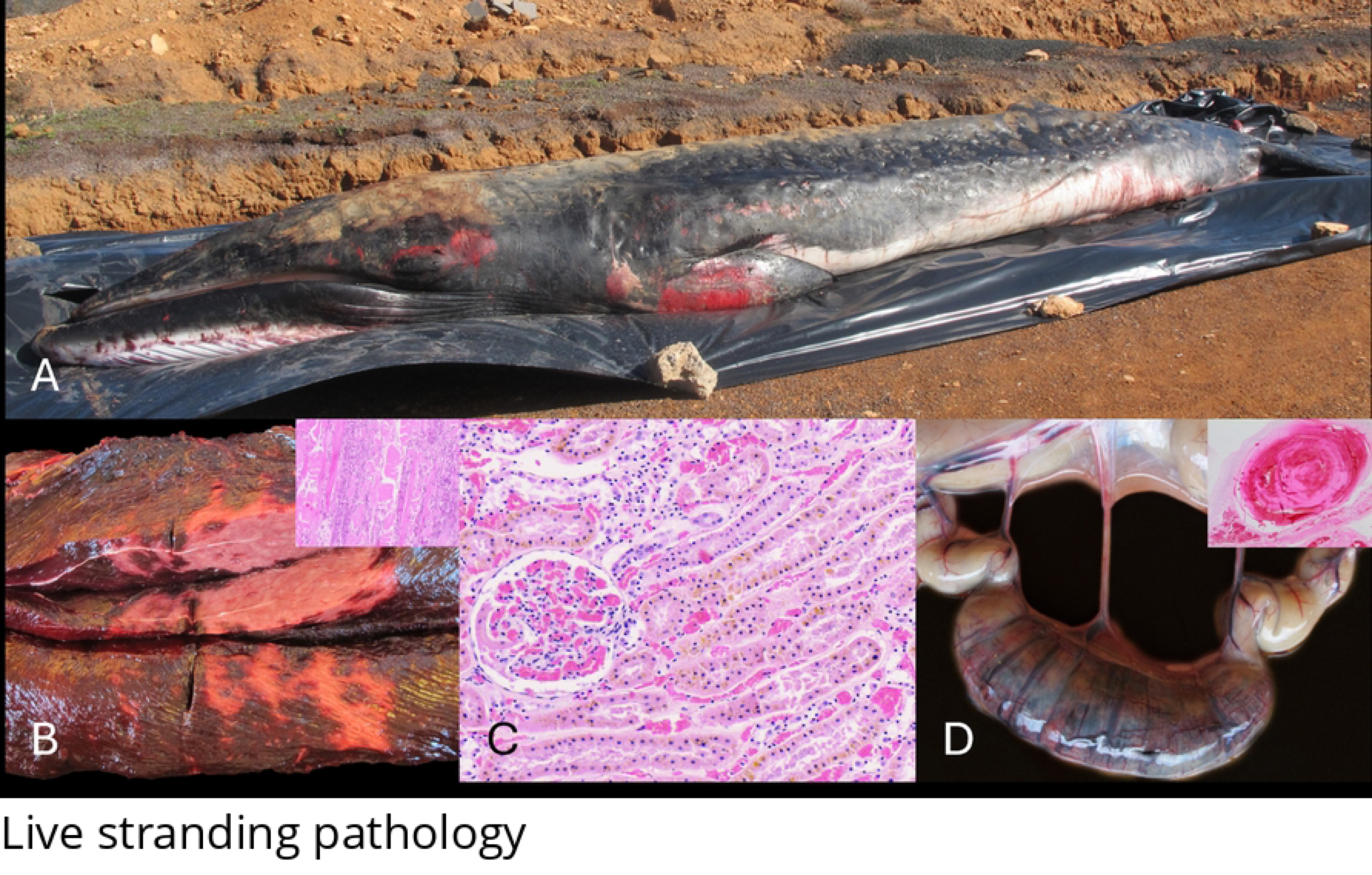
Panel of pathologies associated to live stranding syndrome in cetaceans stranded in the Canary Islands (2013-2018). **A.** Animal on necropsy spot (animal no. 42; *B. edeni*). Note multifocal epidermal lacerations and hemorrhages on abdomen, ventral aspect of pectoral fins, and periocular region. Inset. Detail of subendocardial hemorrhages. **B.** Musculo-skeletal myonecrosis (animal no. 22; *D. delphis)*. The *longissimus dorsi* exhibited multifocal to coalescing, deep, and well delimitated areas of myonecrosis. Inset. The interstitium is expanded by lymphocytes, plasma cells, and macrophages with frequent segmental and floccular necrosis and regeneration signs. **C.** Pigment-associated acute tubular injury (animal no. 180; *G. macrorhynchus*). Multifocal pigment accumulation within the tubuloepithelial cells. **D.** Hemorrhagic enteritis with thrombosis (animal no. 22; *D. delphis*). Segmental jejunal dilation and intraluminal hemorrhage. Inset. Blunt and fused intestinal villi (upper left) with large intravascular thrombi within the lamina propria and abrupt dissection of the lamina muscularis mucosae.

These animals had common underlying infectious inflammatory processes including viral encephalitis (animals no. 43, 84, 180), herpesviral balanoposthitis (animal no. 43), bacterial broncho-interstitial pneumonia and hepatitis (animal no. 18), and bacterial necroulcerative and suppurative gastritis. An adult short-finned pilot whale (animal no. 211) had lesions compatible with uremic syndrome due to urethral obstruction.

A short-beaked dolphin (animal no. 22) exhibited segmental jejunal dilation with dissecting hemorrhage separating the mucosa and muscularis layers with prominent intravascular thrombosis (Fig. 4D).

## 2. Anthropic pathologic categories

### 2.1. Interaction with fishing activities

IFA implicated a total of 17/224 (7.5%) animals belonging to five species (4 odontoceti; 1 mysticeti) with overrepresentation of the Atlantic spotted dolphin and short-beaked common dolphin. Main etiologic diagnoses were sharp trauma by fishing gear (e.g., harpoon, hook, longline; 41.2%), entanglement (29.4%), by-catch (23.5%), blunt trauma (5.9%).

Macroscopic findings included cutaneous net marks (animals no. 11, 113, 115, 122, 145, 153) (Fig. 5A and 5B), fractures (e.g., mandible, maxilla, vertebrae, occipital; animals no. 130, 151, 218), fishing hook perforations (e.g., sublingual, esophageal, oral) (Fig. 5C and 5D) and fishing gear incisions (animals no. 32, 42, 88, 151, 152, 156, 161) (Fig. 5E), intravascular gas dilatations (animals no. 81, 156, 161), pyothorax (animal no. 156), and multicavity hemorrhage. A short-beaked common dolphin (animal no. 130) exhibited a ∼30° deviation of the caudal peduncle, consistent with scoliosis, along with lumbar vertebral fractures and hemorrhage. Most common histologic findings were hemorrhage, segmental myonecrosis, intravascular gas bubbles, bronchial constriction and degenerative myopathic changes were common.

**Fig. 5.**
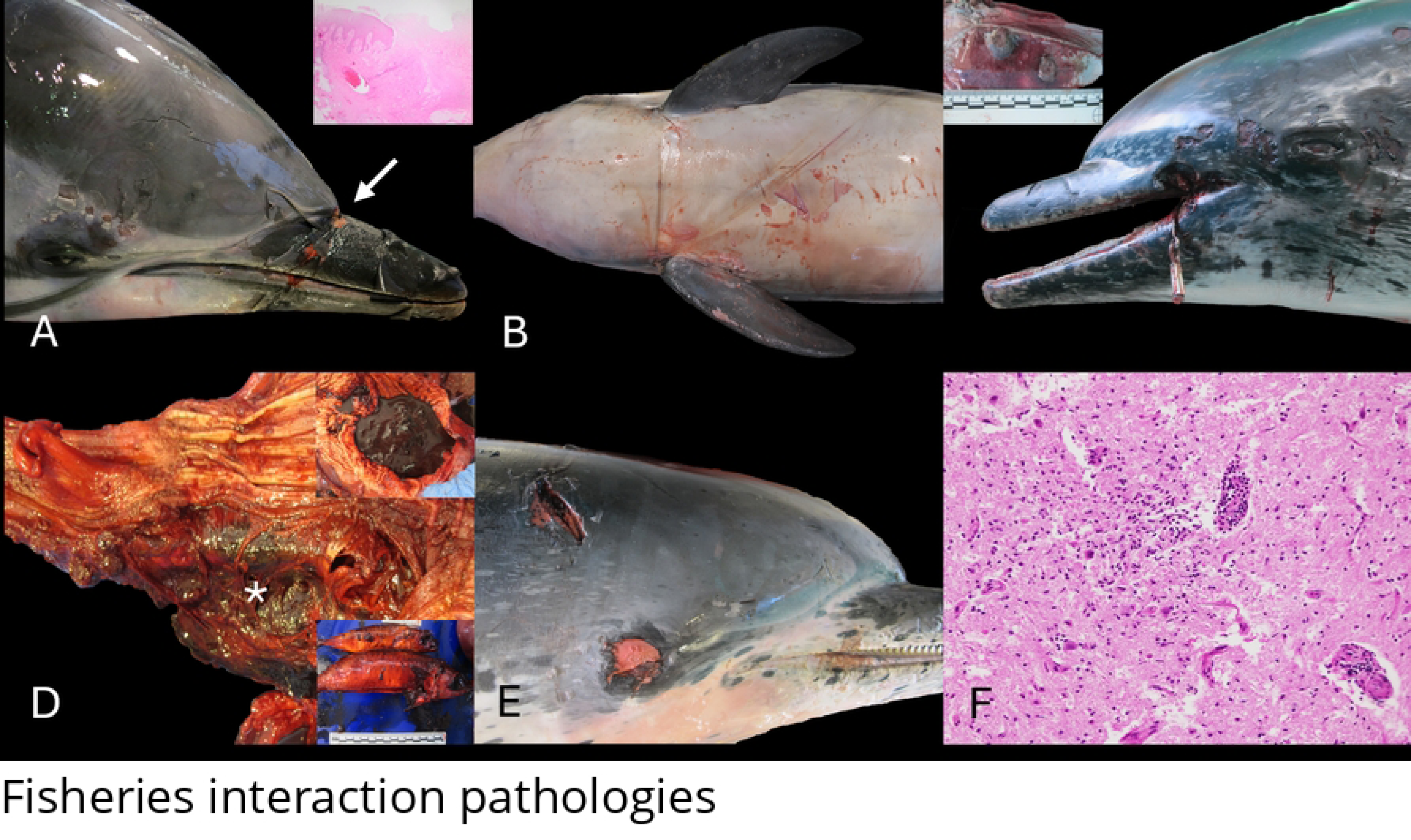
Panel of pathologies associated to interaction with fishing activities in cetaceans stranded in the Canary Islands (2013-2018). **A.** Entanglement with fishing gear (animal no. 153; *S. coeruleoalba*). Circumferential and linear ulcerative dermatitis and stomatitis. Inset. Ulcerative dermatitis with retracted and hyperplastic borders wtih deep reepitelization. **B.** Net marks (animal no. 122; *S. coeruleoalba*). Ventro-Cervical and thoracic zig-zag epidermal rope impressions. **C.** Hook perforation (animal no. 161; *S. frontalis*). Focal perforation and osteomyelitis (not visible) with intralesional large hook. Inset. Multifocal necroulcerative glossitis. **D.** Esophagic perforation (animal no. 88; *S. frontalis*). Oesophageal transmural perforation (asterisk) by a hook (not visible) with tracheal and left hemithorax involvement. Bottom left inset: semi-digested preys. Bottom right inset: keratinized stomach filled with abundant blood **E.** Anthropogenic-associated incision with fishing gear (animal no. 151; *S. frontalis*). Focal incisive left cephalo-cervical wound with sharp edges. **F.** Encephalitis (animal no. 42; *S. frontalis*). Lymphoplasmacytic encephalitis with gliosis and perivascular cuffing’s formation (unknown etiology).

Chronic entanglement was detected in a mink whale calf (animal no. 11) with severe, bilateral, chronic dermatitis and stomatitis in the rostral mandible and necroulcerative glossitis.

Six of 17 (35.3%) animals presented concomitant infectious processes, including: lymphoplasmacytic encephalitis and meningoencephalitis (animals no. 34, 42, 81, 151, 156) (Fig. 5F), lymphohistiocytic myocarditis with necrosis and hemorrhage (animal no. 130), and presumptive herpesviral gastric and biliary infection (animal no. 81) Multiorgan lymphoproliferative disease compatible with a lymphoma was noted in a short-beaked common dolphin (animal no. 219) that succumbed to fishing interaction.

### 2.2. Foreign-body associated pathology

FBAP was identified in 3/224 (1.3%) animals including one Cuvier’s beaked whale (animal no. 5), one pygmy sperm-whale (animal no. 47), and one striped dolphin (animal no. 79). The stripped dolphin was cachectic, and the pygmy sperm whale had a moderate body condition. The advanced decomposition status hampered body condition assessment in the Cuvier’s beaked whale.

Macroscopic lesions included intestinal perforation with intraluminal and intra-abdominal large plastic foreign bodies, compatible with a plastic bag and packaging material (animal no. 5) (Fig. 6A and 6B); ulcerative gastritis with intralesional nematodes and numerous plastic foreign material including wrapping and hard plastics (animal no. 47 and 79) (Fig. 6C). Animal no. 47 had multisystemic parasitism with cardiovascular and renal crassicaudiasis. Incidental or seemingly irrelevant ingestion of foreign bodies was noted in 7 animals (no. 37, 39, 48, 60, 64, 154, 187). Fishing line, plastic bags, female underwear (e.g., bra), nylon segments, or packaging remnants were findings within the gastrointestinal tract [12].

**Fig. 6.**
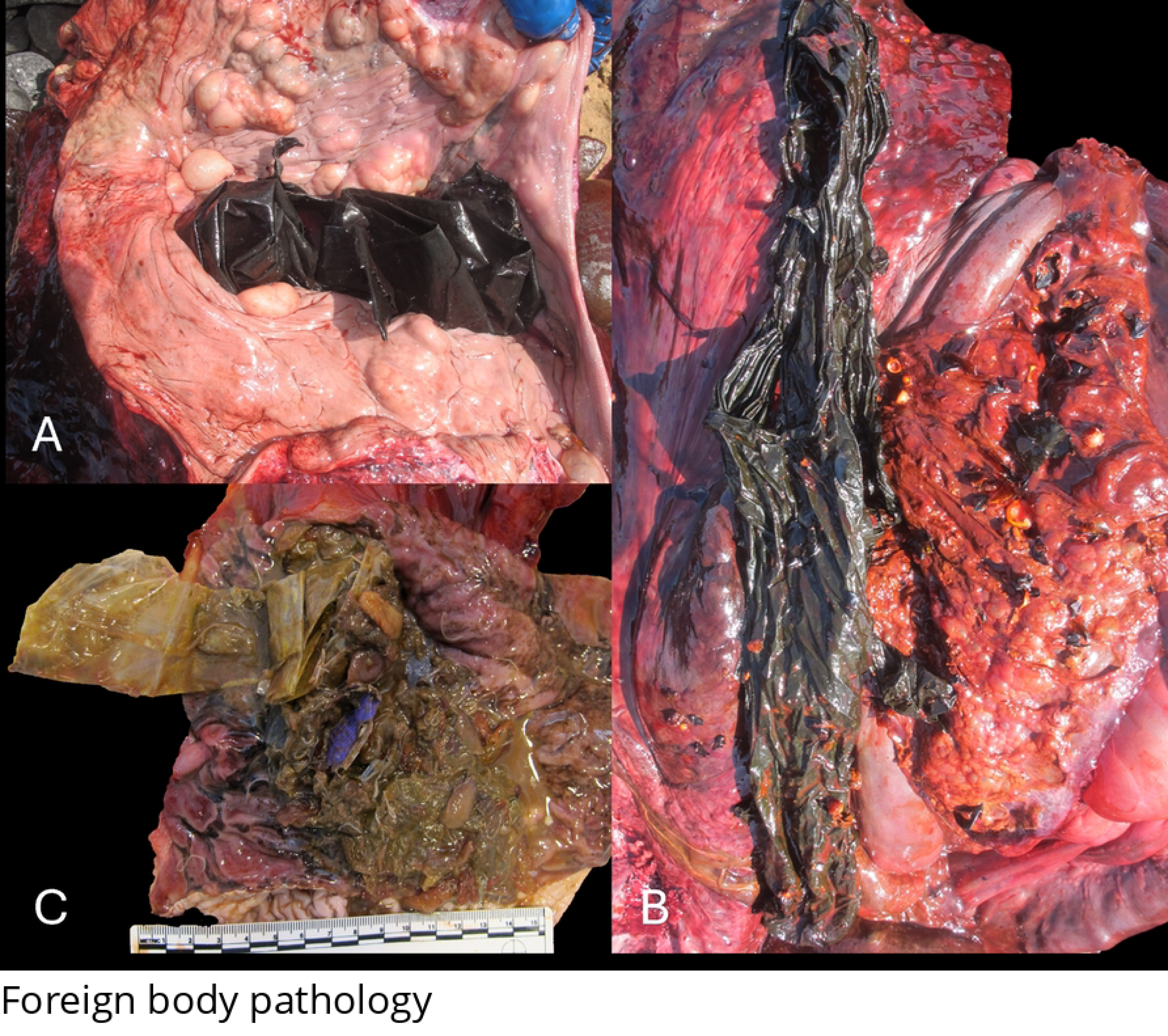
Panel of foreign-body associated pathologies in cetaceans stranded in the Canary Islands (2013-2018). **A.** Plastic foreign body (animal no. 5; *Z. cavirostris*). Wrapped plastic foreign object within the second gastric chamber with hemorrhage within the underlying mucosa (not visible). **B.** Plastic foreign body (animal no. 5; *Z. cavirostris*). Detail of a 1 × 1 m black plastic bag with moderate amount of semi-digested food. **C.** Plastic foreign body (animal no. 47; *K. breviceps*). Multiple plastic foreign objects (blue and transparent) with numerous embedded anisakid nematodes within the glandular stomach.

### 2.3. Vessel collisions (VC)

VC was diagnosed in 9/224 (4%) animals including three sperm whales (animal no. 19, 59, 101), three Cuvier’s beaked whales (animals no. 53, 104, 138), two pygmy sperm-whales (animals no. 656 and 925), and one Bryde’s whale (animal no. 159). Three animals were calf or juveniles and six adults. Macroscopic findings, often consistent with sharp and blunt-force trauma, included incised-contused wound on the dorsolateral left side of the head with fractures of the occipital and parietal bones and hemopericardium (animal no. 19) (Fig. 7A), partial/complete peduncle amputation (animal no. 656 and animal no. 104), fractures of the transverse processes of the lumbar vertebrae (animal no. 12), perpendicular bodily amputation caudal to the dorsal fin with extrusion of the abdominal viscera (animal no. 53), incised-contused wound with vertebral fracture cranial to the dorsal fin (animal no. 59) (Fig. 7B), two penetrating wounds, in the peduncle and abdomen, with protrusion of viscera hemoabdomen, and severance of the spinal cord (animal no. 101), complete peduncle amputation with hemoabdomen, and traces of anti-fouling paint on the cutting edges and sectioned vertebral bodies (animal no. 104), partial section of the aortic trunk, hemothorax and hemopericardium (animal no. 818), incised-contused wound with multiple vertebral fractures cranial to the dorsal fin (animal no. 159) (Fig. 7C), and complete amputation of the peduncle with fractures of the vertebral bodies and ribs (animal no. 214).

**Fig. 7.**
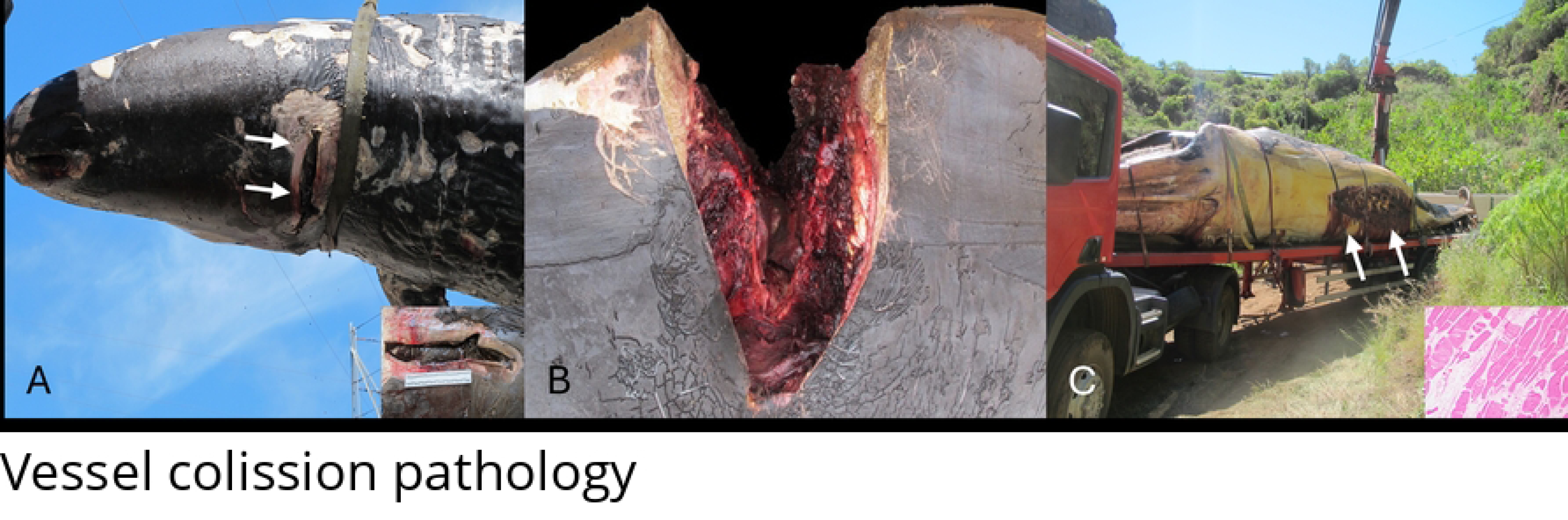
Panel of pathologies associated to vessel collision in cetaceans stranded in the Canary Islands (2013–2018). **A.** Suspended cadaver during transportation (animal no. 19*; P. macrocephalus)*. Sharp-blunt wound (∼50–60 cm) on the left dorsolateral cephalo-cervical region (white arrows). Inset: angular and sharp wound edges. **B.** Sharp-blunt trauma (animal no. 59; *P. macrocephalus*). Wedge-shaped transverse sharp-blunt wound cranial to the dorsal fin with vertebrae fractures. **C.** Cadaver transported by a crane to the necropsy spot (animal no. 159; *B. edeni*). Linear sharp wound (∼1 m) on the dorsum cranial to the dorsal fin with fracture and exposure of dorsal vertebral processes (white arrows). Inset: Segmental myonecrosis with sarcoplasm retraction and floccular pattern (*Longissimus dorsi*)

The main histopathologic findings in these animals were acute myodegeneration and necrosis in the skeletal and cardiac striated muscle. Segmental discoid myodegeneration and necrosis prevailed. Intrapulmonary fat embolism was confirmed by osmium tetroxide technique in five animals (animals no. CET 12, 19, 53, 59, 104) [67,68].

Co-morbidities in these animals were ulcerative gastritis (animal no. 12), verminous arteritis (animals no. CET 53, 104, 138), pyogranulomatous nephritis by *Crassicauda* sp. (animals no. 104 and CET 138), and fibrosing cholangiohepatitis with bile duct hyperplasia (animal no. CET 159).

## 3. Discussion

This study marks the third generation of long-term, pathology-based research on stranded cetaceans in the Canary Islands, reflecting a legacy of tireless multidisciplinary collaboration and generational continuity [2,3]. Despite the inherent difficulties and limitations in the pathologic scrutiny of stranded cetaceans, much knowledge has been gained and this has enabled a firm foundation for a better understanding of the main threats to these species across the Canarian archipelago.

This study assessed 224 of 316 stranded cetaceans, including 18 species (15 odontocetes and 3 mysticete). Previous studies in the archipelago examined 138 of 233 (59%), including 17 species (1999-2005) (2) and 236/ 320 (74%), including 20 species (2006-2012) (3). Comparatively, in this study, we provided pathologic data for two Bryde whales and one killer whale, species not previously assessed in the area. Such variations in the stranding epidemiology may reflect migratory patterns, carcass accessibility, and/or altered ocean dynamics potentially affecting currents, prey distribution, and cetacean migration routes [69,70].

Species and analysis rates from relatively similar but geographically distant studies vary notably: 133/445 (33%) [35], 302/550 (55%) [36], and 222/472 (21%) [24].

We did not detect sex-biased stranding distribution, in agreement with the preceding study [3].Furthermore, we observed high mortality in young individuals with reduction in adult individuals and peaking again in aging animals, aligning with previous observations [71]. Death of young individuals was common in Atlantic spotted dolphin, minke whale, and sperm whale. Díaz-Delgado et al. [3] reported a threefold increase in mortality among calf and juvenile sperm whales, directly associated with vessel collisions. We also detected higher strandings in the eastern Islands, wherein coastline length, assiduity, carcass accessibility, trade winds, currents, topography, and species-specific habitats are known to play a role.

The annual average of strandings during 2013-2018 (53) was higher than between 2006 and 2012 (45.7), and significantly above the average registered between 1995 and 2005 (39). Nonetheless, the stranding peak namely between March and June, remained [2,3,72].

The cause of death was determined in 98.9% of fresh carcasses, 96.4% of carcasses with moderate autolysis, and 64.9% with advanced autolysis, which is in agreement or higher than previous studies (2, 3).

## 1. Natural pathologic categories

Unlike previous studies, this work contemplated cases without loss of nutritional status (NPGNS), cases with significant loss of nutritional status (NPSLNS), or without body condition assessment due to autolysis (NPWNS).

### 1.1. Natural pathology not associated with significant loss of nutritional status

As in previous studies, infectious disease was most prevalent (58.13%), followed by parasitic diseases (27.9%)

CeMV was identified in two short-finned pilot whales and one striped dolphin with typical lesions in the nervous, respiratory and lymphoid systems [16,31,61,73–76]. Unusual lesions in these animals were fibrinonecrotizing laryngitis, lymphoplasmacytic prostatitis, and orchitis with cytoplasmic and intranuclear viral inclusion bodies [77]. Six strains of CeMV have been recognized, representing a major threat for wildlife cetaceans [54,78–82] Prospective IHC and molecular analyses (for CeMV in suspicious cases have revealed higher incidence than initially known for CeMV infection in the archipelago [2,61,83]. A considerable number of animals (12/43; 27,9%) in this category had some degree of CNS inflammation; however, a cause was not readily evident. Ongoing and prospective analyses will help clarify their etiologies, including assessment of emerging pathogens such as a highly pathogenic avian influenza A (H5N1) [29,30,84,85].

Multisystemic herpesvirosis with neurological involvement was detected in one striped dolphin and two Atlantic spotted dolphins. Also, localized cutaneous herpesvirosis was identified in a pygmy sperm whale. Alpha- and gamma-herpesviruses have been reported to cause disease in various cetacean species worldwide, primarily affecting the nervous, integumentary, and urinary systems [61,83,86–93].Arbelo et al. [94] detected an α-herpesvirus in a Cuvier’s beaked whale associated with fibrino-necrotizing vasculitis and coagulative necrosis in the spleen and multiple lymph nodes. Additionally, coinfections with *Brucella* spp. and *Staphylococcus aureus* were detected in both Atlantic spotted dolphins. Herpesvirus–bacteria coinfections are well-documented in cetaceans, including a minke whale with *Brucella*-meningoencephalitis [95] and a striped dolphin with atlanto-occipital osteoarthritis linked to *Mycoplasma* sp. [3]. Neurobrucellosis was detected in a striped dolphin calf.. *Brucella ceti* and *B. pinnipedialis* have been reported in several cetacean species, mainly associated with severe inflammation of the CNS (1, 2, 3). Our findings further corroborate *Brucella* sp. as a main differential etiology for meningoencephalitis cetaceans from Canary Island waters [49], underscoring the need for caution when handling stranded animals due to its zoonotic potential.

Poxvirus was identified via PCR in a Rissos’s dolphin. Cutaneous lesions are well described in published reports [96].

We detected *Erysipela rhusiopathiae* in two Atlantic spotted dolphins and one short-beaked common. All individuals had multiorgan intracytoplasmic bacillary bacteria within macrophages and suppurative and histiocytic myocarditis with necrosis. Similar lesions were noted previously in a bottlenose dolphin and an Atlantic spotted dolphin [52] Although these strandings occurred over several years, they occurred in a relatively delineated geographical location in the coastline of Tenerife. These data are suggestive of a localized area with *E*. *rhusiopathiae* circulation, but the potential source of infection, likely from terrestrial origin, has not been identified. The first report of *E. rhusiopathiae* in a common dolphin from the Canary Islands is provided.

Pulmonary infection by *S. aureus* and herpesvirus coinfection was detected in an Atlantic spotted dolphin. Arbelo et al. [2] identified *S. aureus* as the causative agent of a bronchopneumonia in a Fraser dolphin (*Lagenodelphis hosei*). Infections by *S. aureus* have been widely described in cetaceans, with common respiratory involvement [97–101].

Cerebral mucormycosis with concurrent CeMV infection was detected in a short-finned pilot whale. Concomitant fungal and viral infection of the CNS is unusual [3,73,102]. Other reports of fungal encephalitis in cetaceans include: *Cunninghamella bertholletiae* [103]*, Aspergillus fumigatus* [104]*, Fusarium oxysporum* [105], and *Coccidioides immitis* [106,107].

Severe necroulcerative stomatitis and periostitis by presumptive *Conchoderma auritum* was observed in an Atlantic spotted dolphin. Such lesion severity associated with epibionts is *Conchoderma* is exceedingly rare or undocumented.

We detected *Sarcocystis* spp. in one short-beaked common dolphin, one striped dolphin, one Atlantic spotted dolphin and one Risso’s dolphin. Intrasarcoplasmic cysts were seen in epaxial muscle, diaphragm, tongue, heart, and mammary gland. Inflammatory reaction was none or minimal, aligning with previous studies [108].

Severe pterygoid sinusitis by *Nasitrema* sp. *Crassicauda* sp. and *Stenurus* sp. was seen in two Atlantic spotted dolphins, one striped dolphin, one common dolphin, one Risso’s dolphin, and one bottlenose dolphin. Gross and microscopic findings were consistent with previous reports [2,3], including fatal vestibulocochlear neuritis and meningoencephalitis [63,107,109]. Severe sinusitis may impair the echolocation system, potentially affecting predatory and social behaviors, among other vital neurological functions.

Multiorgan infestation by *Crassicauda* sp. was observed in several animals, consistent with previous studies [3,110,111]. Less prevalent *Crassicauda*-lesions were pyogranulomatous prostatitis with urethral obstruction caused by adult nematodes, and renal vascular and rete mirabile nematode egg embolism in an adult Atlantic spotted dolphin. *Crassicauda* sp. egg vascular embolization has been reported in cetaceans [3,112–114]; in case of high parasitic burden, there might be concern for circulatory disturbances including organ hypoperfusion.

An Atlantic spotted dolphin had multiple renal cysts associated with interstitial nephritis. Similar renal lesions are rarely reported in cetaceans and may be linked to age or genetic factors [115]. In this case, the cystic lesions were likely related to inflammation and distal nephron obstruction. This is the first description of renal cysts in this species and in cetaceans from Canary Island waters.

A uterine leiomyoma associated with postpartum pyometra and uterine prolapse was reported in an Atlantic spotted dolphin [62]. Uterine neoplasms have been documented in multiple instances in cetaceans [116–118]; however, concomitant prolapse is a rare occurrence.

A Blainville’s beaked whale exhibited gross and histologic findings consistent with systemic gas embolism[10,119]. This animal had gastric foreign bodies as noted in aged beaked whales (2) and pilot whales [2]. Currently recognized causes of gas embolism in cetaceans include mid-frequency sonar [10,120], bycatch/ forced ascent [4,5,121], ship collision, and behavioral, mainly on deep divers [122], and failed predatory attempts [123]; however, in a number of cases, the exact etiopathogenesis remains undetermined after reasonable exclusion of the former [10,122,123]. Our data suggest advanced age is a common factor and may be a predisposing cause in these cases.

### 1.2. Natural pathology associated with significant loss of nutritional status

Infectious (61.1%) and parasitic etiologies (30.5%) were the most prevalent, which is in agreement with a previous study [3]. CeMV and concurrent herpesviral infection was detected in one striped dolphin, one short-finned pilot whale and one common dolphin. In addition, *Vibrio* sp., a known bacterial pathogen in cetaceans [124–128], was isolated from the lungs in the short-finned pilot whale. (106–110). The latter represents a unique case of triple coinfection by CeMV, HV and *Vibrio* sp. in an odontocete from the Canarian waters. Furthermore, herpesvirus was identified in seven individuals belonging to five odontocete species. Histological findings were consistent with previous reports, with particularly severe involvement of the nervous and respiratory systems. Notably, a Blainville’s beaked whale with herpesviral detection in kidney and lung had concurrent scapulohumeral and atlanto-occipital osteoarthritis, and lymphoplasmacytic bronchointerstitial pneumonia associated with *Brucella* spp. and *P. damselae* susp. *damselae*, respectively. Infectious arthritis by *Brucella* sp. has been reported in multiple cetaceans [49,129,130]. *P. damselae* was recently described causing pneumonia in bottlenose dolphins [131]. To our knowledge, this is a unique case involving a triple bacterial and viral coinfection in a Blainville’s beaked whale. In addition, severe suppurative bronchopneumonia caused by *P. damselae* was diagnosed in an rough-toothed dolphin dam and its calf. These results underscore the prevalence and pathogenic potential of *P. damselae* in Canarian waters [3].

Bacterial culture identified *Clostridium perfringens* in two pygmy sperm whales with fibrinous polyserositis, splenomegaly, hepatitis, lymphadenitis and vasculitis with intralesional bacilli. Septicemia by *C. perfringens* has been described in multiple cetaceans [3,77,132–134]. These cases represent the first description of *C. Perfringens* infection in pygmy sperm whales. These animals also had findings consistent with previous reports of dilated cardiomyopathy. The latter could have potentially contributed to morbidity and mortality in these cases by decreased cardiovascular resilience.

(86,111,117). A juvenile Atlantic spotted dolphin had pyogranulomatous tracheitis, thyroiditis, and laryngitis by *Rhizopus arrhizus* [64].

A Blainville beaked-whale had severe pyogranulomatous encephalitis with intralesional adult *Nasitrema delphini* [63]. Typical inflammatory foci in the auditory and central nervous system recapitulated features consistent with aberrant parasite migration through the inner ear and brain seen in other cetacean species[109,135,136].

A short-finned pilot whale presented severe urethral obstruction with intraluminal adult *Crassicauda* sp. nematodes, as well as pyelonephritis, nephrolithiasis, tracheitis, laryngitis, and ulcerative glossitis with thrombosis. Altogether, the lesions observed were highly suggestive of uremic syndrome, which is a condition poorly documented in cetaceans. Blood chemistry analyses were not available in this case.

Right pneumothorax following rupture of the right tracheal bronchus due to severe obliterative bronchial nematodiasis was seen in short beaked common dolphin. Cases of pneumothorax in cetaceans have been previously documented [123,137–139]; recently, a broader term “airway leak syndrome” which would include pneumothorax, has been suggested in marine mammals [140]. Other possible causes of bronchial rupture where reasonably ruled out in this case.

Lumbar kypho-lordosis associated with hemivertebrae was seen in an Atlantic spotted dolphin calf. Congenital vertebral column malformations have been described in various cetacean species [141,142].

Polycystic gas hepatopathy” recapitulating features of Budd-Chiari-like syndrome was diagnosed in a striped dolphin. The presence of hepatic gas cysts has been associated with pre-existing liver disorders (e.g., hepatobiliary trematodiasis) exacerbated by gas bubble formation [143].

A cavernous hemangioma in the mesenteric lymph node was described in a short-beaked common dolphin. Similar proliferative vascular lesions in the lung have been reported in short-beaked common dolphins and Atlantic bottlenose dolphins, possibly linked to parasitism [144].

### 1.3. Natural pathology without assessable body condition

Parasitic and infectious etiologies prevailed in this category. Six Cuvier’s beaked whales had moderate to severe verminous arteritis and nephritis by *Crassicauda* spp., [20,145]. The severity of renal and vascular disease appeared to increase with age. In these cases, vascular disease may have a significant impact in foraging dives and other vital actions resulting in compromised health status. In a subset of these animals (animals 60, 61, and 103), *Crassicauda anthonyi* was identified based on morphological features and mitochondrial *cox1* and ITS-2 sequencing [60].

In this category, CeMV caused lymphoplasmacytic meningoencephalitis in a striped dolphin. Another striped dolphin had atlanto-occipital arthritis of likely bacterial origin. Fibrinosuppurative balanoposthitis of likely bacterial was noted in a sperm whale calf. The exact causes for the last two cases could not be determined.

An Atlantic spotted dolphin had a uterine laceration and polyserositis following a dystocic parturition. The fetus presented prominent macrosomia and subcutaneous edema. Uterine rupture is a relatively rare occurrence in cetaceans [3,146].

Based on gross and microscopic pathologic findings, presumptive metabolic steatohepatitis was diagnosed in a striped dolphin [147,148]. Blood tests to assess secondary hepatic conditions were unavailable.

### 1.4. Neonatal/ perinatal pathology

NPP encompasses a broad spectrum of conditions affecting individual viability in utero, and/or shortly after birth (2,3). In agreement with previous observations, fetal distress was a common finding, followed by dystocia, premature maternal-social separation and/or neonatal weakness, intra-interspecific trauma, and infectious disease processes [2,3]. Detailed lung evaluation is essential to assess fetal stress or perinatal complications in cetaceans [2,3,149–152].

Suppurative omphalitis with cardiac hemorrhage was seen in a Bryde whale calf, recapitulating features noted in pygmy sperm whale calf [3]. Although bacterial infection is suspected, the causative agent remained unidentified. Notably, we report the only known case of a stranded killer whale calf in the Canary Islands. This animal had pulmonary hemorrhage, edema, keratin squames and numerous coccobacilli within alveoli and bronchioles. *Vibrio furnissii* was isolated from the spleen. Previously detected in bottlenose dolphins [153] and known to cause disease in humans [154], *V. furnissii* may have contributed to septicemia in this case.

### 1.5. Traumatic intra- interspecific interaction

IITIs are commonly reported in cetacean species and seem to be overrepresented in short-finned pilot whales; the factors triggering them are variegated [42,155–161]. We observed ITIIs comprised primarily short-finned pilot whales and pygmy and dwarf sperm whales (*Kogia sima*). In agreement with previous studies, we observed multiorgan (e.g., pulmonary) and cavitary hemorrhage (hemoabdomen, hemothorax), bone fractures, partially digested ingesta and rake marks. Typical skeletal myonecrosis with discoid degeneration patterns and pigmentary tubulonephrosis were common. Notable cases were a short-finned pilot whale with severe cranioencephalic trauma and extensive cerebral hemorrhage and gas embolism. The (98,100,105) first description of gas embolism in a short-finned pilot whale in the Canary Islands is documented. Additionally, a Sowerby’s beaked whale was diagnosed with multiorgan myofiber embolism resulting from trauma [66].

Fatal killer whale interactions were diagnosed in five animals from four species [161]. Killer whale predatory or playful encounters have been reported in numerous cetacean species [162–166]. These encounters occurred at the time killer whales have historically been recorded in the Canary Islands following seasonal prey movements [161] (Puig-Lozano et al., 2020). Co-morbidities were common in the affected animals.

Asphyxia due to laryngeal luxation associated with one elongated fish (compatible with conger) was diagnosed in a bottlenose dolphin. The fish extended from the pharynx to the cardias. A fatal prey-predator interaction was determined. Similar cases of asphyxia related to prey ingestion have been reported in other species [3,123,167].

### 1.6. Live-stranding stress and/or capture myopathy-related pathology

Live stranding events in cetaceans may induce an extreme stress response in cetaceans [3,168,169]. In such events, homeostatic derangement is accelerated by heat retention, solar radiation, neurogenic shock, and worsening of pre-existing conditions [2,3]. Live-stranded cetaceans often die during or after rescue (e.g., handling) due to muscle damage resembling capture myopathy [170–174]. Live-stranding-related pathology was the main factor for death in 18 individuals (8%); striped and short-beaked common dolphins were overrepresented yet at an overall lower rate than previous studies in the archipelago [2][3]. Lesions mainly involving the integumentary and musculoskeletal system were common, as previously reported [3,152,168,169,173,175,176]. Intracytoplasmic eosinophilic globules within hepatocytes were frequent [177]. Serum markers (e.g., troponin I and C, creatine kinase) have been linked to the stranding and/or rescue process in cetacean species [152,176]. Notable cases were a short-beaked common dolphin with extensive myonecrosis and hemorrhage resembling muscular clostridiosis by *Clostridium chauvoei* or *Cl. septicum*. The exact etiology in these cases remains undetermined.

## 2. Anthropic pathologic categories

### 2.1. Interaction with fishing activities

Identifying lesion patterns linked to interactions with fisheries is key to estimate the impact of this major anthropogenic threat to cetaceans [4,5,178,179]. Distinguishing accidental capture, chronic entanglement, and direct aggression is essential [4,5,178]. We detected evidence of fatal interaction with fishing activities in 17/224 (7.5%) animals. Although several studies have assessed and delineated typical gross and microscopic findings seen in this pathologic category [4,5], this anthropogenic impact remains largely underestimated. Historical and ongoing pressures and extractive activities in oceanic and riverine environments may not always be readily evident upon pathologic examinations [180–183]. Between 2000 and 2005, and 2006 and 2012, an incidence of 19/138 (13.8%) [2] and 10/236 (4.2%) [3], respectively, was detected. Typical macroscopic lesions in these animals included cutaneous net marks, bone fractures, perforations, and intracavitary hemorrhage (2,3). A short-beaked common dolphin showed marked scoliosis at the peduncle, along with multiple vertebral fractures. Traumatic scoliosis has been previously reported [3]. Histologically, multiorgan hemorrhage, as well as segmental myonecrosis were common. Four animals had incised wounds and internal organ lacerations caused by fishery gear (e.g., harpoons, hooks). Six animals had evidence of an underlying infectious process (e.g., meningoencephalitis, myocarditis). A short-beaked common dolphin had a lymphoproliferative disease process resembling a hepatosplenic lymphoma [184].

### 2.2. Foreign body associated pathology

In recent decades, excessive plastic use has become a major threat to wildlife, particularly marine species, ecosystems, and human health. Plastic is involved in 92% of marine debris interactions across nearly 700 species [185]. Entanglement and ingestion affect many cetaceans [12,186], with rising reports of plastic ingestion in 48 species [187]. Additionally, microplastic impacts on food webs are an emerging concern [188–190]; the effects of microplastics on cetaceans remain unknown. Young age and deep-diving behavior have been proposed as predisposing factors for foreign body ingestion [2,3,12,191].

In this study, death was attributed to foreign body ingestion in a juvenile Cuvier’s beaked whale, a pygmy sperm whale calf, and a striped dolphin calf. These animals had intestinal perforation with and ulcerative gastritis with plastic fragments and packaging debris. Moreover, plastic ingestion was considered an incidental finding in seven animals classified in other pathologic entities.

Arbelo et al. [2] reported foreign body ingestion in two Atlantic spotted dolphins, a Cuvier’s beaked whale, and a striped dolphin. Díaz-Delgado et al. [3]reported gastric obstructions, gastric perforation, and chronic entanglement in Cuvier’s beaked whale, a pilot whale, a juvenile Gervais’ beaked whale, and a minke whale. A detailed review on foreign body ingestion and the pathologic findings associated during the study period has been published [12].

Neurological impairment (e.g., inflammation) could promote plastic ingestion [2]. Furthermore, poor body condition is significantly associated with plastic debris ingestion [12], and deep-diving species are more affected than shallow divers [12,192]. Our data, along with previous reports, show a high representation of deep-diving species e.g., *Z. cavirostris*, *M. europaeus*, *G. griseus*, *K. breviceps*, affected by this pathology, supporting this hypothesis.

### 2.3. Vessel collisions

A VC implies the impact between any part of a boat (typically the hull or propeller) and a marine mammal [193]. Such collisions often result in severe or fatal trauma [67,178]. In general, whale strike incidence peaked between 1950 and 1980, coinciding with the global shipping boom [7]. By 2005, the international whaling commission (IWC) prioritized ship strikes and created the “Ship Strikes Working Group” [194]. A detailed review of marine mammal collisions is provided by Schoeman et al. [195].

VC can cause sharp-force trauma (from propellers) and/or blunt-force trauma (from non-rotating parts like the hull), leading to abrasions, contusions, lacerations, and fractures [67,178,196]. In very decomposed cadavers, such injuries may be difficult to assess [178,196].

We detected 9 fatal vessel collision events, mainly affecting deep-diving species e.g., sperm whales, Cuvier’s beaked whales, pygmy sperm whales, and Bryde’s whales. The species distribution in our study, including odontocetes and mysticetes, is similar to previous studies in the archipelago [2,3]; however, we recorded a lower number of sperm whales dead to VC [3]. From 23 sperm whales stranded (3.8 per year), only 11 were fully assessed. From the remaining 12, at least three, had evidence of possible vessel collision; however, advanced decomposition and logistical difficulties hindered full pathologic assessment.

The animals in this category had typical macroscopic and histologic findings[2,3,7,9,67,178] pulmonary fat embolism was confirmed in five animals by osmium tetroxide (OsO₄) histochemical technique [68].

In the Canary Islands, a growing number of cargo ships and inter-island ferries, operating at speeds from 15 to over 30 knots, overlap spatially and seasonally with peaks on sperm whale populations (spring and summer) [67,197,198]. Sperm whales are listed as vulnerable by the IUCN [199] and are the species most affected by VCs in the region [2,3,67].

Female sperm whales and their offspring may be more vulnerable to vessel collisions [200], particularly young whales with slower swimming and longer surface times [7,9,201]. Whitehead and Shin (2022) postulated the annual collision rate to be similar to the population growth rate, suggesting vessel strikes could neutralize or severely limit the recovery of local sperm whale populations over time.

In alignment with previous studies [2] [3], most collisions occurred along Tenerife’s eastern coast, a key corridor with high maritime traffic, dominated by fast and ultra-fast ferries [9,67,200].

## Conclusion

Our results indicate that 15% of cetacean deaths were attributable to anthropic activity, while natural pathologies accounted for 85% in the Canary Islands between 2013 and 2018. Natural an anthropic pathologic entities, in decreasing order, were: natural pathology associated with good nutritional status (19.2%), natural pathology associated with significant loss of nutritional status (16%), intra- and interspecific traumatic interactions (13.4%), natural pathology without stablished body condition (8.5%), live-stranding stress syndrome (8%), interaction with fishing activities (7.6%), vessel collision (4%), foreign body-associated pathology (1,34%). The CD could not be established in (13.4%) individuals. Infectious and parasitic events were the most prevalent within natural pathologic entities. This study consolidates health monitoring efforts of stranded cetaceans in the Canary Islands by compiling valuable post-mortem data across a wide range of species, including newly described pathologic conditions, and could serve as a useful guidance for veterinarians, biologists and caregivers as well as a valuable resource for ongoing and future conservation initiatives.

## Acknowledge

This work is dedicated to all those who, faithfully and soulfully, rely on science to shed light and stand up for the truth. All authors would like to thank the incommensurate effort of laboratory technicians, members and volunteers of The Cetacean Stranding network and associated non-governmental stranding associations: Society for the Study of Cetaceans in the Canary Archipelago (SECAC; Vidal Martin) and Canarias Conservación, with special mention to Manuel Carrillo.

